# Rapid, sensitive, full genome sequencing of Severe Acute Respiratory Syndrome Virus Coronavirus 2 (SARS-CoV-2)

**DOI:** 10.1101/2020.04.22.055897

**Authors:** Clinton R Paden, Ying Tao, Krista Queen, Jing Zhang, Yan Li, Anna Uehara, Suxiang Tong

**Author notes:** Address for correspondence: Suxiang Tong, Division of Viral Diseases, Centers for Diseases Control and Prevention, 1600 Clifton Rd NE, Mailstop H18-6, Atlanta, GA 30329. These first authors contributed equally to this manuscript (Clinton R Paden, Ying Tao).

## Abstract

SARS-CoV-2 recently emerged, resulting a global pandemic. Rapid genomic information is critical to understanding transmission and pathogenesis. Here, we describe validated protocols for generating high-quality full-length genomes from primary samples. The first employs multiplex RT-PCR followed by MinION or MiSeq sequencing. The second uses singleplex, nested RT-PCR and Sanger sequencing.

---

In December 2019, SARS-CoV-2, the etiological agent of Coronavirus Disease 2019 (COVID-19), emerged in Wuhan, China. Since then it has rapidly spread to the rest of the world (1-3). As of April 16, 2020, there have been 1,991,562 confirmed cases, including 130,885 deaths, in 185 countries or regions (4).

Initial sequencing of SARS-CoV-2 showed limited genetic variation between cases, but did document specific changes that may be useful for understanding the source and transmission chains (5-8). Because SARS-CoV-2 has shown the capacity to spread rapidly and lead to a range of presentations in infected persons, from asymptomatic infection to mild, severe, or fatal disease, it is important to identify genetic variants in order to understand any changes in transmissibility, tropism, and pathogenicity. Sequence data can be used to inform decisions to better manage the spread of disease.

In this report, we describe the design and use of two PCR-based methods for sequencing SARS-CoV-2 clinical specimens. The first is a multiplex PCR panel followed by sequencing on either the Oxford Nanopore MinION or Illumina MiSeq. When coupled with MinION sequencing, the protocol can be implemented outside a traditional laboratory and can be completed in a single workday, similar to previous mobile genomic surveillance of Ebola and Zika virus outbreaks (9, 10). Additionally, we provide a complementary singleplex, nested PCR strategy, which improves sensitivity for samples with lower viral load and is compatible with Sanger sequencing.

## The Study

On January 10, 2020, the first SARS-CoV-2 genome sequence was released online (11). That day, we designed two complementary panels of primers to amplify the virus genome for sequencing. For one panel, we used the PRIMAL primer design tool (9) to design multiplex PCRs to amplify the genome in using only a few PCR reactions (Appendix). The final design consists of 6 pools of primers, targeting amplicon sizes of 550 base pairs (bp) with 100bp overlaps, to allow for sequencing on either the ONT MinION or Illumina MiSeq. For the second panel, we designed sets of primers to generate nested, tiling amplicons across the SARS-CoV-2 genome (Appendix), for enhancing sensitivity in samples with lower viral loads. Each amplicon is 322-1030bp with an average overlap of 80bp. They are designed to be amplified and sequenced individually on Sanger instruments but may also be pooled for sequencing on next-generation sequencing platforms.

To determine the sensitivity of each sequencing strategy, we generated a set of six ten-fold serial dilutions of a SARS-CoV-2 isolate (12). Virus RNA was diluted into a constant background of A549 human cell line total nucleic acid (RNaseP CT 29). We quantitated each dilution using the CDC SARS-CoV-2 rRT-PCR for the N2 gene (13) (data not shown). The six dilutions spanned CT values from 22-37, corresponding to ca. 2 to 1.8 × 10^5^ copies. We amplified triplicate samples at each dilution using the multiplex PCR pools. Next, we pooled, barcoded, and made libraries from each sample’s amplicons using the ligation-based kit and PCR barcode expansion kit (methods in Appendix). MinION sequencing was performed on an R9.4.1 or R10.3 flow cell until we obtained >1-2M raw reads. From those, 50-60% of reads could be demultiplexed (data not shown). Additionally, we sequenced these amplicons using the Illumina MiSeq for comparison (methods in Appendix).

For MinION sequencing, the reads were basecalled and analyzed using an in-house read mapping pipeline (detailed in Appendix). For samples with CT ≤ 29, we obtained >99% SARS-CoV-2 reads and >99% genome coverage at 20X depth, decreasing to an average of 93% genome coverage at CT 33.2 and 48% at CT 35 (Figure 1A and 1B). Further, we were able to obtain full >20X genomes within the first 40-60 minutes of sequencing (Figure 1C).

**Figure 1.**
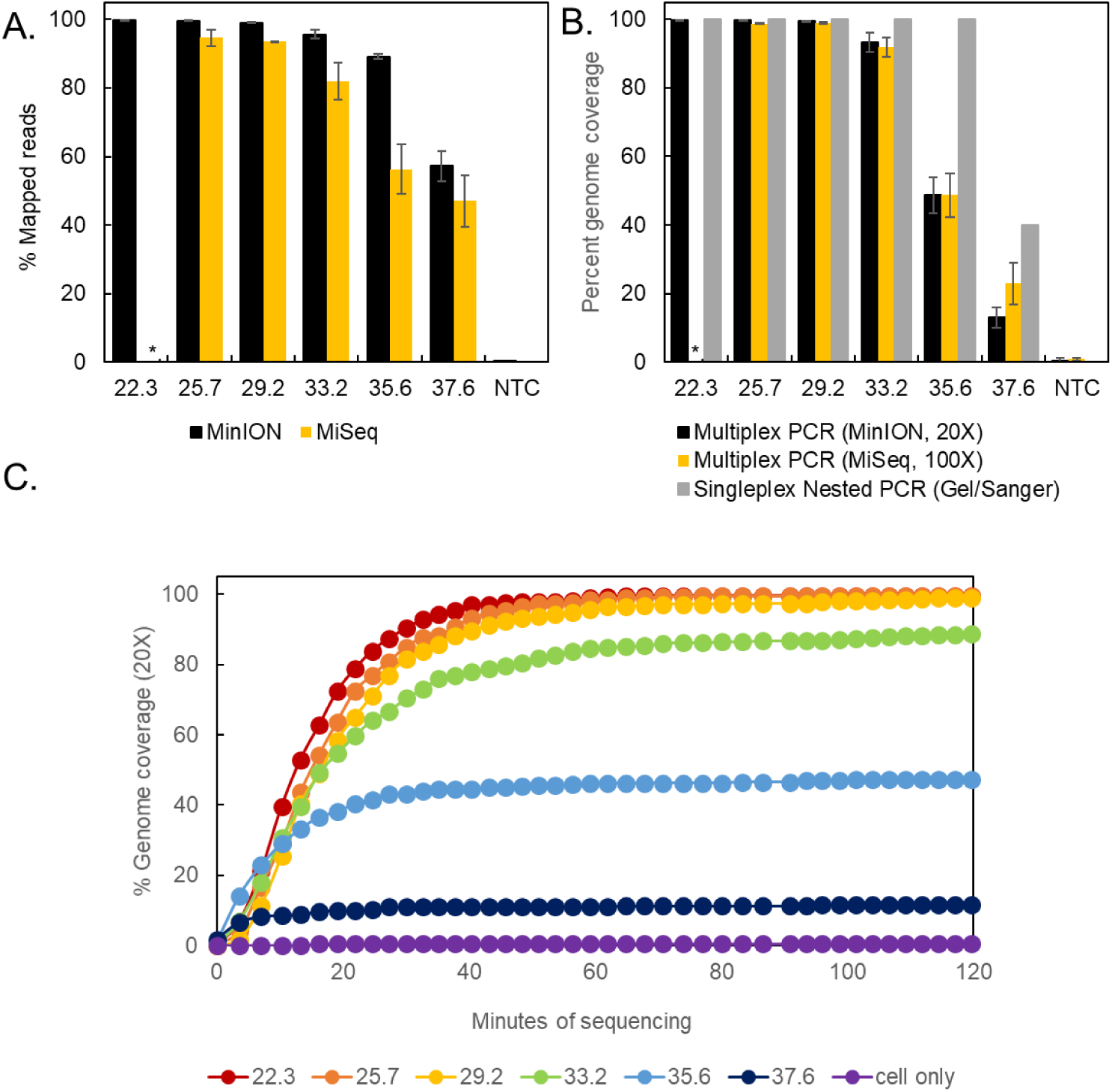
Limits of detection. Triplicate serial dilutions of SARS-CoV-2 isolate A12 (12) amplified using the singleplex or multiplex primer set. The multiplex amplicons were barcoded, library-prepped, and sequenced on a MinION or MiSeq. (A) Percent of reads that map to the virus genome for each sample. (B) Percent of virus genome that is covered at >20X depth by the multiplex amplicons on the MinION (black) or >100X depth on the MiSeq (orange), or covered by the nested, singleplex amplicons (grey) (measured by presence or absence on a gel). (C) Real-time analysis of MinION sequencing data. Each data point represents the average 20X genome coverage of three replicates.

Consensus accuracy, including SNPs and indels, is critical for determining coronavirus lineage and transmission networks. For high consensus level accuracy, we filtered reads based on length, mapped them to the reference sequence (RefSeq NC_045512), trimmed primers based on position, and called variants with Medaka (https://github.com/nanoporetech/medaka) (details in Appendix). Each Medaka variant was filtered by coverage depth (>20X) and by the Medaka model-derived variant quality (>40). Here we used the variant quality score as a heuristic to filter remaining noise from the Medaka variants, compared to Sanger-derived sequences. After these steps, the data approaches 100% consensus accuracy (Table 1). Identical results were found using the R9.4.1 pore, through the CT 33.2 samples (data not shown). We noted larger deletions in some of the CT 33.2+ samples which likely reflect biases from limited copy numbers.

**Table 1.** Genome consensus accuracy

| **Virus tier (C<sub>T</sub>)** | **%Coverage (20X)<sup>a</sup>** | **Indels** | **Indel bases** | **SNPs** | **%Identity<sup>b</sup>** |
| --- | --- | --- | --- | --- | --- |
| 22.3 | 99.659 | 0 | 0 | 0 | 100 |
|  | 99.722 | 0 | 0 | 0 | 100 |
|  | 99.635 | 0 | 0 | 0 | 100 |
| 25.7 | 99.635 | 0 | 0 | 0 | 100 |
|  | 99.615 | 0 | 0 | 0 | 100 |
|  | 99.642 | 0 | 0 | 0 | 100 |
| 29.2 | 99.508 | 0 | 0 | 0 | 100 |
|  | 99.635 | 0 | 0 | 0 | 100 |
|  | 99.615 | 0 | 0 | 0 | 100 |
| 33.2 | 93.024 | 1 | 1 | 0 | 100 |
|  | 93.603 | 2 | 35 | 0 | 100 |
|  | 87.894 | 0 | 0 | 0 | 100 |
| 35.6 | 41.653 | 1 | 1 | 0 | 100 |
|  | 51.266 | 0 | 0 | 1 | 99.993 |
|  | 50.821 | 1 | 15 | 2 | 99.987 |
| 37.6 | 14.634 | 0 | 0 | 1 | 99.977 |
|  | 9.317 | 0 | 0 | 0 | 100 |
|  | 12.363 | 0 | 0 | 0 | 100 |

<sup>a</sup> The 5' and 3' ends are primer sequence, so 100% coverage is not possible
<sup>b</sup> Percent of covered bases identical to reference sequence, excludes indels and low-coverage bases

In the MiSeq data, we observed a similar trend in percent genome coverage at 100X depth, and a slightly lower percent mapped reads, compared to Nanopore data (Figure 1A and B). Increased read depth using the MiSeq potentially allows increased sample throughput, however the number of available dual unique barcodes limits actual throughput.

For the nested, singleplex PCR panel, we amplified the same serial dilutions with each nested primer set (methods in Appendix). The endpoint dilution for full genome coverage is approximately CT 35 (Figure 1B). At the CT 37 dilution, we observed significant amplicon dropout—at this dilution, there are <10 copies of the genome on average per reaction.

These protocols enabled rapid sequencing of the initial clinical cases of SARS-CoV-2 in the United States. For these cases, we amplified the virus genome using the singleplex PCR amplicons, sequencing them with both MinION and Sanger instruments to validate MinION consensus accuracy. The MinION produced full-length genomes in <20 minutes of sequencing, while Sanger data was available the following day.

We used the multiplex PCR strategy in subsequent SARS-CoV-2 clinical cases (n=167), ranging in CT values from 15.7 to 40 (mean 28.8, median 29.1). In cases below CT 33, we observed an average of 99.02% specific reads and 99.2% genome coverage at >20X depth (Figures 2A and 2B). Between CT 30-33, genome coverage varied by sample, and declined dramatically at higher CT values, analogous to the isolate validation data. For these samples, we multiplexed 20-40 barcoded samples per flowcell. Enough data is obtained with 60 minutes of MinION sequencing for most samples, though for higher titer samples 10-20 minutes of sequencing is sufficient (Figure 2C).

**Figure 2.**
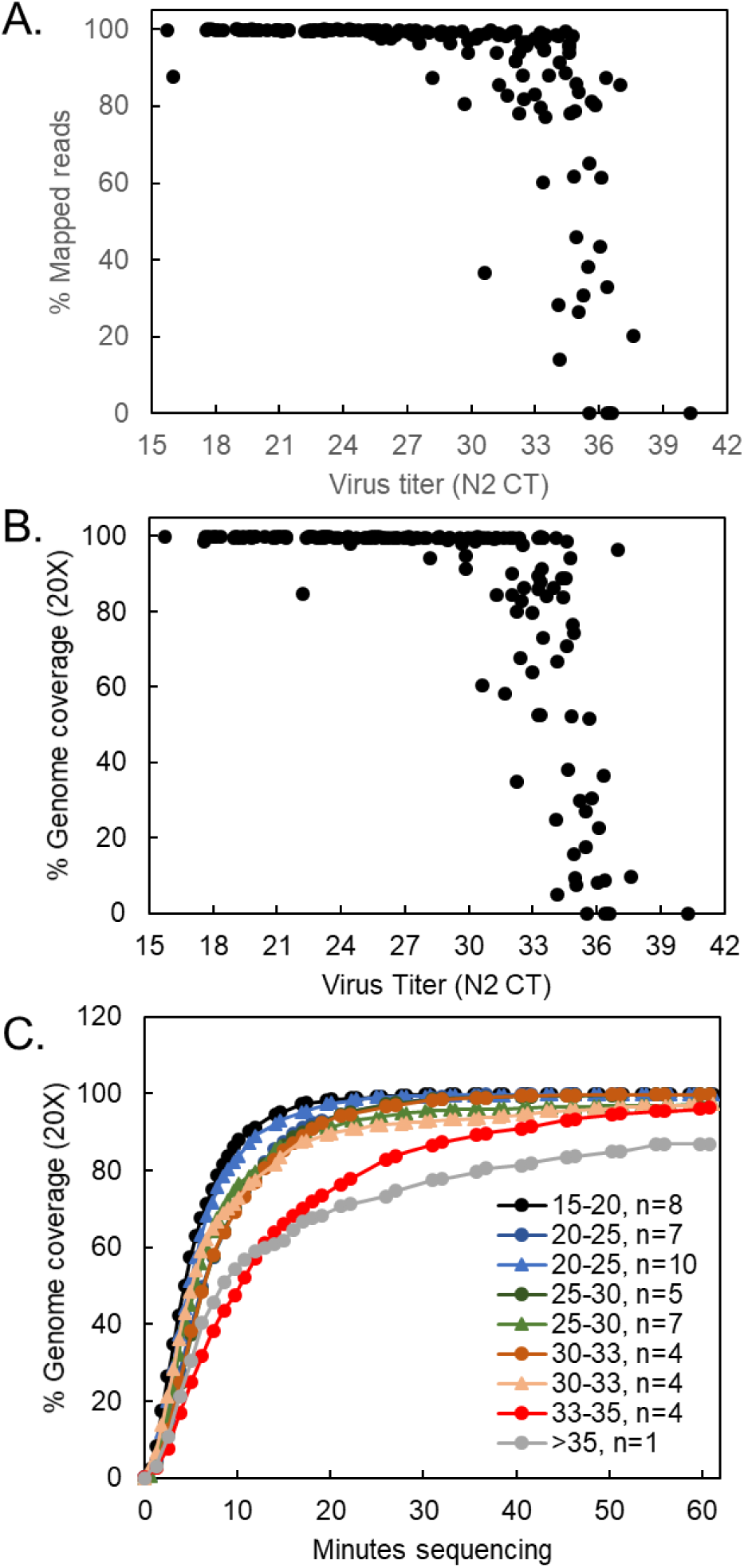
Sequencing SARS-CoV-2 clinical samples. (A) Percent mapped and (B) percent genome coverage for 167 clinical SARS-CoV-2 samples, amplified with multiplex PCR strategy and sequenced on the MinION. (C) Time-lapse of 20X genome coverage obtained by MinION for clinical specimens at the indicated CT values. Data points represent the average coverage for the indicated number of samples

Up-to-date primer sequences, protocols, and analysis scripts are found at https://github.com/CDCgov/SARS-CoV-2_Sequencing/tree/master/protocols/CDC-Comprehensive. Data from this study is deposited in the NBCI SRA (BioProjects PRJNA622817 and PRJNA610248).

## Conclusions

Full genome sequencing is an indispensable tool in understanding emerging viruses. Here we present two validated PCR target-enrichment strategies that can be used with MinION, MiSeq, and Sanger platforms for sequencing SARS-CoV-2 clinical specimens. This ensures that most labs have access to one or more strategies.

The multiplex PCR strategy is effective at generating full genome sequences up to CT 33. The singleplex, nested PCR is effective up to CT 35, varying based on sample quality. The turnaround time for the multiplex PCR MinION protocol is about 8 hours from nucleic acid to consensus sequence, compared to Sanger sequencing at about 14-18 hours (Table 2). Importantly, the multiplex PCR protocols offer an efficient, cost-effective, scalable system, and add little time and complexity as sample numbers increase (Table 2). The results from this study suggest multiplex PCR may be used effectively for routine sequencing, complemented by singleplex, nested PCR for low virus-titer samples and confirmation sequencing.

**Table 2.**
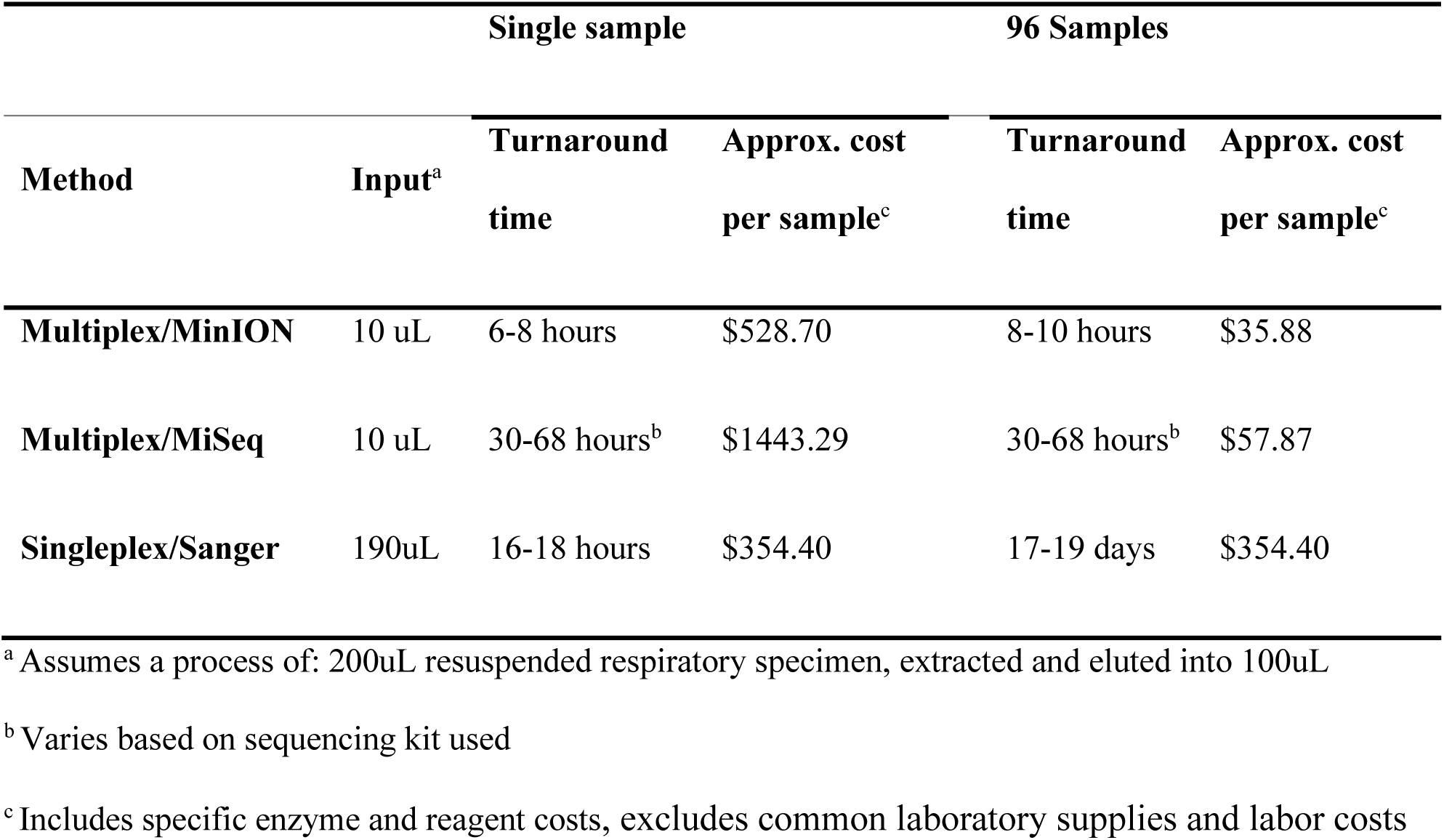
Comparison of input, time, and cost requirements for sequencing one or 96 specimens

## Supporting information

Appendix - Sequencing Protocol

## Acknowledgments

We would like to acknowledge the efforts of those in the Respiratory Viruses Branch at CDC who helped in organizing samples for this study, including Azaibi Tamin, Jennifer Harcourt, Natalie Thornburg, Shifaq Kamili, Xiaoyan Lu, and Stephen Lindstrom.

## Disclaimers

The findings and conclusions in this report are those of the authors and do not necessarily represent the official position of the Centers for Disease Control and Prevention.

## Author Bio

(first author only, unless there are only 2 authors)

Clinton Paden is a virologist and bioinformatician in the CDC Pathogen Discovery Team, within the Division of Viral Diseases. His work is in identifying and characterizing novel and emerging pathogens.

