## Appendix - Sequencing Protocol for "Rapid, sensitive, full genome sequencing of Severe Acute Respiratory Syndrome Virus Coronavirus 2 (SARS-CoV-2)"

### Protocols for SARS-CoV-2 sequencing

**Pathogen Discovery Team**
**NCIRD/DVD/RVB**
**Centers for Diseases Control and Prevention**

### Disclaimers

The findings and conclusions in this report have not been formally disseminated by the Centers for Disease Control and Prevention and should not be construed to represent any agency determination or policy.

The protocols described here are for research purposes only and should not be used in place of approved diagnostic testing.

### Singleplex nested RT-PCR

#### Protocol Notes

To complete this protocol, 185 uL of extracted template is needed. For samples between Ct 27 and 35, two rounds of nested RT-PCR are recommended; for samples up to Ct 27, one round of RT-PCR is recommended. The resulting PCR products can be individually proceeded with Sanger sequencing, or they can be pooled for Oxford Nanopore or Illumina sequencing, depending on the number of samples and availability of sequencing platforms.

**See Appendix C for recommended plate setups.**

#### Required Reagents

| Company | Product | Catalog number |
|---|---|---|
| Thermo (Invitrogen) | Superscript III one-step RT-PCR with Platinum Taq High Fidelity DNA polymerase | 12574035 |
| Sigma Aldrich (Roche) | Protector RNase inhibitor | 3335402001 |
| Takara | LA Taq DNA polymerase with GC buffer | RR02AG |
| | Nuclease Free water | |
| | 50 uM Primers | |

#### Procedure

##### 1. First round of RT-PCR

1.1. Prepare the first-round master mix as below. Please note, the protocol is generic as all 38 primer pairs require the same master mix (see Appendix A). For each SARS-CoV-2 sample to be sequenced, 38 individual PCR reactions are required.

| Component | Volume (uL) |
|---|---|
| Water | 1.75 |
| 2x Buffer (2.4mM $MgSO_4$) | 12.5 |
| 5mM $MgSO_4$ | 4.5 |
| 50uM Primer For | 0.25 |
| 50uM Primer Rev | 0.25 |
| RNase Inhib. 40U/uL | 0.25 |
| SSIII / Platinum Taq high fidelity | 0.5 |
| Pre-mix | 20 |
| Template (RNA) | 5 |
| Total | 25 |

1.2. Add 5uL of RNA template to each of the 38 PCR reactions. Spin tubes/plates down and proceed to PCR.

1.3. Perform first round PCR with the cycling parameters as below.
60°C 1min, decrease 0.5°/sec, 94°C, 2 min; 40 cycles of 94°C 15 seconds, 55°C 15 seconds, 72°C 60 seconds; 72°C 7 minutes, 12°C

#### 2. Second round of semi-nested or nested PCR

2.1. After first round RT-PCR is complete, prepare the master mix for $2^{nd}$ round of semi-nested or nested PCR as below. Please note, the protocol is generic as all 38 second round primer pairs require the same master mix. Primer information is located in Appendix A. For the $2^{nd}$ round of semi-nested- or nested PCR, there are 38 individual PCR reactions for each sample to be sequenced.

| Component | Volume (uL) |
|---|---|
| Water | 5.75 |
| 2× GCBuffer I | 12.5 |
| dNTP Mixture (2.5 mM each) | 4 |
| 50uM Primer For | 0.25 |
| 50uM Primer Rev | 0.25 |
| TaKaRa LA Taq™ (5 units/μl) | 0.25 |
| Pre-mix | 23 |
| Template (1R product) | 2 |
| Total | 25 |

2.2. Add 2 uL of the corresponding first round PCR product to the second round PCR master mix. Spin tubes/plates down and proceed to PCR.

2.3. Perform second round PCR with the cycling parameters as below.

94°C, 3 min; 40 cycles of 94°C 15 seconds, 55°C 15 seconds, 72°C 60 seconds; 72°C 7 minutes, 12°C

2.4. Following the completion of second round PCR, run 3 uL of all 38 PCR reactions on 1% agarose gels or fragment analyzer to check for amplification.

### Sanger sequencing

#### Required Reagents

| Company | Product | Catalog number |
|---|---|---|
| Thermo (Applied Biosystems) | ExoSap-It | 78201.1.ML |
| Thermo (Applied Biosystems) | BigDye v3.1 cycle sequencing kit | 4337455 |
| Princeton Separations | Centri-sep 96 well plates | CS-963 |
| | Nuclease Free water | |
| | 5 uM Primers | |

#### Procedure

1. Transfer 10 uL of each PCR reaction to new tubes/plate for ExoSap cleanup. Add 4 uL ExoSap-It to each PCR reaction (10 uL) and incubate at 37°C for 15 minutes, followed by 80°C for 15 minutes on a thermocycler.
2. Prepare sequencing master mix as below.
   Sequencing primers for each amplicon are listed in Appendix B

| Component | Volume (uL) |
|---|---|
| Water | 5.5 |
| 2x Buffer | 2 |
| 5uM Primer | 1 |
| BigDye 3.1 enzyme | 1 |
| Pre-mix | 9.5 |
| Template (2R PCR product) | 0.5 |
| Total | 10 |

3. Add 0.5 uL of corresponding ExoSap cleaned PCR product to each sequencing reaction mix. Spin tubes/plates down and proceed to sequencing PCR.
4. Perform sequencing PCR with the parameters listed below:
   96°C, 2 min, 30 cycles of 96°C 30 seconds, 50°C 15 seconds, 60°C 3 mins, 4°C forever
5. Following sequencing PCR, clean-up of sequencing reactions is performed with Centri-Sep 96-well plates following the manufacturer's instructions (Appendix G) with one addition. 20 uL nuclease free water is added to the 96-well collection plate prior to the final spin.
6. The 96-well collection plate with the cleaned sequencing sample plus water is loaded onto the ABI sequencer.
7. Sequencher 5.4 is used for data analysis of Sanger PCR data.

### Multiplex PCR

#### Protocol Notes

This protocol uses 10 uL of template for each sample. The pooled, multiplexed PCR products can be followed with nanopore sequencing or Illumina MiSeq sequencing depending on the number of samples and available sequencing platforms. We have been able to sequence full genomes reliably under Ct 30, and depending on the sample, up to Ct 33.

This protocol was adapted from Quick J et al. *Nat Protoc*. 2017 Jun;12(6):1261-1276.

#### Required reagents

| Company | Product | Catalog number |
|---|---|---|
| Thermo Fisher (Invitrogen) | SuperScript IV 1st strand synthesis system | 18091200 |
| NEB | NEBNext Q5 Hot Start HiFi PCR Master Mix | M0543L |
| | Nuclease Free water | |
| | Primers | |

#### Procedure

##### 1. Generate primer pools

1.1. Prepare primers as 50 uM primer stocks.

1.2. Add an equal volume of each 50 uM primer stock to six 1.5mL Eppendorf tubes labeled as pool 1, 2, 3, 4, 5, and 6. Primers for each pool are listed in Appendix D.

1.3. Prepare 10 uM working concentration by diluting each pool 1:5 with nuclease free water.

##### 2. First-strand synthesis

2.1. Mix the following components.

| Component | Volume(uL) |
|---|---|
| RNA (template) | 10 |
| Random primer 25uM | 2 |
| dNTPs | 1 |
| Total | 13 |

2.2. Denature the template-primer-dNTP mix at 65°C for 5 minutes.

2.3. Place on ice for 5 minutes.

2.4. Add the following components to the template-primer-dNTP mix:

| Component | Volume (uL) |
|---|---|
| 5x SSIV buffer | 4 |
| 0.1 M DTT | 1 |
| RNAse inhibitor | 1 |
| SSIV RT (200 units/uL) | 1 |
| Total | 20 |

2.5. Incubate in a thermal cycler at the following temperatures:
25°C 10 minutes, 50°C for 10 minutes, 85°C for 10 minutes, hold at 4°C.

2.6. Spin down. Can be stored at -20°C

2.7. Add 1 uL RNase H and incubate at 37°C for 20 minutes

#### 3. Multiplex PCR

3.1. Mix the following components in 6 wells of a PCR plate or strip tube.

| Component | Volume (uL) |
|---|---|
| NEBNext Q5 Hot Start HiFi PCR Master Mix | 15 |
| PCR grade water | 10.2 |
| Primer pool 1, 2, 3, 4, 5, or 6 (10uM) | 1.8 |
| Total | 27 |

3.2. Add 3 uL of cDNA from above to each tube.

3.3. Run the following PCR program:
98°C 30 seconds, 40 cycles of 98°C 15 seconds, 65°C 5 minutes.
Note: fewer cycles may be used, but 40 cycles is used to maximize detection of lower-titer samples.

3.4. Optional: Run a 2% agarose gel for each multiplexed PCR reaction pool 1, 2, 3, 4, 5, and 6 to check for specific bands of the correct size (0.4-0.6 kb).

3.5. Pool 20 uL from each of 6 tubes of multiplexed PCR reactions in a 0.3 mL tube in a PCR strip or a well in PCR plate (the total volume is 120 uL).

3.6. Add 1X ratio (120 uL) of AMPure XP beads to the PCR product pools.

3.7. Purify according to standard AMPure protocol (see Appendix E).

3.8. Elute in 80 uL water.

3.9. Quantitate 1 uL of cleaned PCR products using Qubit dsDNA HS kit (Appendix F).

3.10. Optional: Run a 2% agarose gel and load 3 uL of cleaned PCR products to check for specific bands of the correct size (0.4-0.6 kb).

### Nanopore Sequencing

#### Protocol Notes

This protocol takes advantage of the multiplexing density afforded by the "PCR Barcoding Expansion 1-96" kit. This protocol is derived from Oxford Nanopore's protocols available at http://community.nanoporetech.com.

Required reagents for Nanopore barcoding and sequencing:

| Company | Product | Catalog number |
|---|---|---|
| NEB | NEBNext Ultra II End-repair/dA tailing module | E7546 |
| NEB | Blunt/TA ligase master mix | M0367 |
| NEB | NEBNext Quick Ligation Module | E6056 |
| TaKaRa | TaKaRa LA Taq DNA Polymerase with GC Buffer | RR02AG |
| Beckman Coulter | Agencourt Ampure XP Beads | A63880/A63881 |
| Oxford Nanopore Technologies | Nanopore Ligation Sequencing Kit (1D) | SQK-LSK109 |
| Oxford Nanopore Technologies | PCR Barcoding Expansion 1-96 | EXP-PBC096 |
| Oxford Nanopore Technologies | SpotON Flow Cell (R9.4.1) | FLO-MIN106D |
| Oxford Nanopore Technologies | MinION | MinION Mk1B |

#### Procedure

##### 1. Barcode amplicons

1.1. Mix the following components:

| Component | Volume (uL) |
|---|---|
| 500 ng amplicon DNA | 25 |
| Ultra II end-prep reaction buffer | 3.5 |
| Ultra II end-prep enzyme mix | 1.5 |
| Total | 30 |

1.2. Incubate at 20°C for 10 minutes, 65°C for 5 minutes, hold at 4°C.
1.3. Add 1X ratio (30 uL) AMPure XP beads.
1.4. Purify according to standard AMPure protocol (Appendex E).
1.5. Elute the DNA target from the beads with 17 uL water.
1.6. Optional: quantitate 1 uL of cleaned end-prep DNA using Qubit dsDNA HS kit (Appendex F)
1.7. Mix the following components:

| Component | Volume (uL) |
|---|---|
| Cleaned end-prep DNA | 15 |
| Barcode Adapter | 10 |
| Blunt/TA ligase master mix | 25 |
| Total | 50 |

1.8. Incubate at 20°C for 10 minutes.
1.9. Add 1X ratio (50 uL) AMPure XP beads.
1.10. Purify according to standard AMPure protocol (Appendix E).
1.11. Elute the DNA in 12 uL water.

1.12. Transfer eluate into new PCR plate or well
1.13. Quantitate 1 uL of ligated DNA according to the protocol (Appendix F).
1.14. Mix the following components:

| Component | Volume(uL) |
|---|---:|
| 30ng adapter-ligated DNA | x |
| PCR Barcode primer (one of BC1-BC96) | 1 |
| 2x GC Buffer I | 25 |
| dNTP mix (10mM) | 8 |
| TaKaRa LA Taq (5U/uL) | 0.5 |
| Water | 50 – x |
| Total | 50 |

1.15. Mix by pipetting and spin down
1.16. Run the following PCR program:
95°C 3 minutes; 18 cycles of 95°C for 15 seconds, 62°C for 15 seconds, and 72°C for 1 minute; final extension 72°C 7 minutes; hold at 4°C.
1.17. Add 1X ratio (50 uL) of AMPure XP beads.
1.18. Purify according to standard AMPure protocol (Appendix E).
1.19. Elute the DNA target from the beads with 25 uL water.
1.20. Quantitate 1 uL cleaned, barcoded PCR products with Qubit dsDNA HS kit (Appendix F).

#### 2. Prepare Nanopore Ligation-based Library

2.21. Pool the barcoded PCR products equally by mass.
2.22. Prepare LSK109 ligation-based libraries by mixing the following components:

| Component | Volume(uL) |
|---|---:|
| 1 ug pooled barcoded sample | x |
| DNA CS | 1 |
| Ultra II End-prep reaction buffer | 7 |
| Ultra II End-prep enzyme mix | 3 |
| Water | 49-x |
| Total | 60 |

2.23. Incubate at 20°C for 10 minutes, 65°C for 5 minutes, hold at 4°C.
2.24. Add 1X ratio (60 uL) of AMPure XP beads.
2.25. Purify according to standard AMPure protocol (Appendix E).
2.26. Elute the DNA target from the beads with 62 uL water.
2.27. To ligate sequencing adapters, mixing the following components:

| Component | Volume(uL) |
|---|---:|
| End-repaired DNA from previous step | 60 |
| Ligation buffer (LNB) | 25 |
| NEBNext Quick T4 DNA Ligase | 10 |
| Adapter Mix (AMX) | 5 |
| Total | 100 |

2.28. Incubate 10 minutes at 20°C
2.29. Add 0.8X ratio (80 uL) of AMPure XP beads
2.30. Purify according to standard AMPure protocol (Appendix E).

2.31. Elute the DNA target from the beads with 15 uL water
2.32. Quantitate 1 uL clean, prepared library with Qubit dsDNA HS kit (Appendix F).

#### 3. Load MinION and sequence

3.33. Set up the MinION flow cell and host computer, including MinKNOW software.
3.34. Open the MinKNOW GUI from the desktop icon and establish a local connection.
3.35. Inset flow cell into MinION.
3.36. Click "Check Flow Cells" at the bottom of the screen then click "Start test." Check the number of active pores available. When the check is complete, it is reported in the Notification panel. Check to ensure it has enough pores for a good sequencing run (warranty for flow cells: 800 nanopores or above checked within 5 days of receipt).
3.37. Thaw the Sequencing Buffer (SQB), Loading Beads (LB), Flush Tether (FLT) and one tube of Flush Buffer (FB) at room temperature before placing the tubes on ice.
3.38. Thoroughly mix the Sequencing Buffer (SQB) and Flush Buffer (FB) tubes by vortexing,
3.39. Spin down the Flush Tether (FLT) tube, mix by pipetting, and return to ice.
3.40. Open the lid of the nanopore sequencing device and slide the flow cell's priming port cover clockwise90 degrees. (The following steps are demonstrated at https://youtu.be/CC11Jlydqrc)
3.41. Set a P1000 pipette to 200 uL, insert the tip into the priming port, turn the wheel until the dial shows 220-230 uL, or until you can see a small volume of buffer entering the pipette tip. Do not remove more than this.
3.42. Visually check that there is continuous buffer from the priming port across the sensor array.
3.43. Prepare the flow cell priming mix: add 30 uL of thawed and mixed Flush Tether (FLT) directly to the tube of thawed and mixed Flush Buffer (FB), and mix by pipetting up and down.
3.44. Load 800 uL of the priming mix into the flow cell via the priming port, avoiding the introduction of air bubbles.
3.45. Wait for 5 minutes.
3.46. Thoroughly mix the contents of the Loading Beads (LB) by pipetting.
3.47. Prepare library for loading my mixing:

| Component | Volume (uL) |
|---|---|
| Sequencing Buffer (SQB) | 37.5 |
| Loading Beads (LB), mixed immediately before use | 25.5 |
| 150-200 ng DNA Library | 12 |
| Total | 75 |

3.48. Gently lift the SpotON sample port cover to make the SpotON sample port accessible.
3.49. Load 200 µl of the priming mix into the flow cell via the priming port (not the SpotON sample port), avoiding the introduction of air bubbles.
3.50. Mix the prepared library gently by pipetting up and down just prior to loading.
3.51. Add 75 uL of sample to the flow cell via the SpotON sample port in a dropwise fashion. Ensure each drop flows into the port before adding the next.
3.52. Gently replace the SpotON sample port cover, making sure the bung enters the SpotON port, close the priming port and replace the MinION lid.
3.53. Start the sequencing run using the MinKNOW software.

#### 4. Generate consensus sequences from MinION data

There are many considerations for generating high-quality consensus data from the MinION. Here are some suggestions for basecalling based on our experience.

Software:

| Software | Source URL |
| --- | --- |
| Guppy 3.4.1+ | https://community.nanoporetech.com/downloads |
| Medaka 0.11.5 | https://github.com/nanoporetech/medaka |
| Minimap2 2.17 (r941) | https://github.com/lh3/minimap2 |
| SAMtools 1.9 | http://www.htslib.org/ |
| BCFtools 1.9 | http://www.htslib.org/ |
| BAMClipper | https://github.com/tommyau/bamclipper |
| cutadapt 2.3+ | https://github.com/marcelm/cutadapt |
| vcf_mask_lowcoverage.pl | https://github.com/CDCgov/SARS-CoV-2_Sequencing |
| IGV | http://software.broadinstitute.org/software/igv/ |

Example commands below have user-supplied variable names bold. You will need to customize the details to your environment.

##### 4.1. Basecalling

Basecalling may also be done using MinKNOW software. If so, you may skip the Guppy basecalling step.

```
# Run Guppy
guppy_basecaller --input_path $rundir --save_path $outputdir -r \
    --config na_r9.4.1_450bps_hac.cfg --barcode_kits EXP-PBC096 \
    --trim_barcodes --require_barcodes_both_ends
# Combine all the output fastq files
mkdir $outputdir/fastq
find $outputdir -name "*.fastq" |while read infile; do
      if [[ $i =~ barcode|unclassified  ]]; then
         outfile=$(grep -Eo "barcode..|unclassified" <<< $infile).fastq
         outfile="fastq/$outfile"
         cat $infile >> $outfile
      fi
done
```

##### 4.2. Filter on quality and length

Filtering out low quality sequence, as well as unexpectedly long and short reads helps tremendously on off-target mapping affecting consensus quality.

```
cutadapt -j $threads -m 300 -M 1200  -q 15 -o $fastqfiltered $fastqfile
```

##### 4.3. Mapping

Download reference sequence from GenBank: MN908947.3

```
minimap2 -L -a -x map-ont -t 12 MN908947.fasta $ fastqfiltered > $samfile
samtools view -b $samfile | samtools sort - -o $bamfile
samtools index $bamfile
```

4.4. Clip primers

This step requires a BEDPE file describing the positions of the primers. It is available at https://github.com/CDCgov/SARS-CoV-2_Sequencing

BAMClipper by default will output at file with the suffix "primerclipped.bam."

Clipping by position allows only primers near the beginning of a read to be trimmed (rather than genuine sequence in the middle of a read), and it is faster than sequence-based trimming (e.g. Porechop).

```
cd $outputdir
bamclipper.sh -b $bamfile -p SC2_200324.bedpe -n 12 -u 80 -d 80
```

4.5. Generate VCF and consensus sequences

Medaka is very lenient with calling variants. We generally require a variant quality score of >= 40 and depth of coverage >= 20 to call a variant. Below 20X coverage, we call an 'N.'

The script to automate the filtering is available at https://github.com/CDCgov/SARS-CoV-2_Sequencing

```
# Generate Medaka VCF File
medaka consensus --model r941_min_high_g344 --threads 12 \ $primerclippedbamfile
$primerclippedbamfile.hdf
medaka variant MN908947.fasta $primerclippedbamfile.hdf $vcf
# Filter variants and generate consensus sequence
vcf_mask_lowcoverage.pl --bam $primerclippedbamfile  \
   --reference MN908947.fasta --vcf $vcf --consout $consensusfasta \
   --depth 20 --qual 40
```

#### 5. Quality control and analysis suggestions

5.6. Watch out for 1-base insertions/deletions. Though consensus calling has improved considerably, there are residual errors. There are several stretches in SARS-CoV-2 that have homopolymers long enough to be problematic

5.7. Do not ignore other deletions. There have been several deletions reported (3, 9, 15, 34bp, 384bp, etc), so keep in mind the difference between a potential real indel and missing amplicon or nanopore error.

5.8. IGV can be useful for examining the "believability" of variants However, some of these 1-2bp indels appear in the reads, but they cannot be confirmed by Illumina or Sanger sequencing. These are either unlucky PCR bias or systematic sequencing error.

### Illumina Library Preparation and Sequencing

#### Protocol Notes

Starting Material: 100 pg–250 ng DNA. We recommend that the DNA be in 1X TE (10 mM Tris pH 8.0, 1 mM EDTA), however, 10 mM Tris pH 7.5–8, low EDTA TE or water are also acceptable. If the input DNA is less than 26 µl, add TE (provided) to a final volume of 26 µl. This protocol is adapted from the NEBNext Ultra II FS protocol, which can be found in its entirety at http://www.neb.com.

For sizing, other devices, such as the 2100 BioAnalyzer, 5200 FragmentAnalyzer, QIAxcel, or LabChipGX may also be used. These vary in quantitation accuracy, so fluorometric quantitation with Qubit (or similar instrument) or qPCR is recommended.

#### Required Reagents

| Company | Product | Catalog number |
|---|---|---|
| New England Biolabs (NEB) | NEBNext Ultra II FS DNA Library Prep Kit for Illumina | E7805S/E7805L |
| New England Biolabs (NEB) | NEBNext® Multiplex Oligos for Illumina (96 Unique Dual Index Primer Pairs) | E6440S/E6440L |
| Beckman Coulter | Agencourt Ampure XP Beads | A63880/A63881 |
| | 10mM Tris-HCl, pH 8.0 | |
| | Molecular biology grade ethanol | |
| | Nuclease-free water | |
| Agilent | High Sensitivity D1000 screen tape | 5067-5584 |
| Agilent | High Sensitivity D1000 reagents | 5067-5585 |

#### Procedure for Library Preparation

##### 1. Fragmentation and End Repair

1.1. Ensure that the Ultra II FS Reaction Buffer is completely thawed. If a precipitate is seen in the buffer, pipette up and down several times to break it up, and quickly vortex to mix. Place on ice until use.

1.2. Vortex the Ultra II FS Enzyme Mix 5-8 seconds prior to use and place on ice.

1.3. Add the following components to a 0.2 ml thin wall PCR tube on ice

| Component | Volume (uL) |
|---|---|
| NEBNext Ultra II FS Enzyme Mix (yellow tube) | 2 |
| NEBNext Ultra II FS Reaction Buffer (yellow tube) | 7 |
| DNA (pooled PCR amplicons) | 26 |
| Total | 35 |

1.4. Vortex the reaction for 5 seconds and briefly spin down. Place in a thermocycler with the heated lid set to ≥75°C and run the following program:

37°C for 7 minutes, 65°C for 30 minutes, 4°C hold indefinitely

#### 2. Adapter Ligation

2.1. Determine dilution for adapter if necessary, see table below. Dilute the NEBNext Adapter for Illumina (red tube) in 10 mM Tris-HCl, pH 8.0 with 10 mM NaCl as indicated below.

| Input DNA in the End Prep reaction | Adapter dilution (volume of adapter: total volume) | Working adapter concentration |
|---|---|---|
| 250 ng - 101 ng | No dilution | 15 uM |
| 100 ng – 5 ng | 10-fold (1:10) | 1.5 uM |
| Less than 5 ng | 25-fold (1:25) | 0.6 uM |

2.2. Add the following components directly to the FS reaction mixture from 1.1(35 uL):

| Component | Volume (uL) |
|---|---|
| NEBNext Ultra II Ligation Master Mix (red tube) | 30 |
| NEBNext Ultra II Ligation enhancer (green tube) | 1 |
| NEBNext adapter for Illumina | 2.5 |

Notes:
- Mix the Ultra II Ligation Master Mix by pipetting up and down several times prior to adding to the reaction.
- The Ligation master mix and ligation enhancer can be mixed ahead of time and is stable for at least 8 hours at 4°C. Do not premix the adapter prior to use in the adapter ligation step.
- The NEBNext adapter is provided in NEBNext Multiplex Oligos for Illumina (96 Unique Dual Index Primer Pairs)

2.3. Set a pipet to 50 uL and pipet entire volume up and down at least 10 times to mix thoroughly. Perform a quick spin to collect all liquid from the sides of the tube.
Note: The NEBNext Ultra II Ligation master mix is very viscous. Care should be taken to ensure adequate mixing of the ligation reaction as incomplete mixing will result in reduced ligation efficiency. The presence of a small amount of bubbles will not interfere with performance.
2.4. Incubate at 20°C for 15 minutes in a thermocycler with the heated lid off.
2.5. Add 3 uL of USER enzyme (red tube) to the ligation mixture.
Note: This step is only required for use with NEBNext adapters. USER enzyme is provided in NEBNext Multiplex Oligos for Illumina (96 Unique Dual Index Primer Pairs)
2.6. Mix well and incubate at 37°C for 15 minutes in a thermocycler with the heated lid set to ≥47°C
2.7. Add 57uL (0.8X) re-suspended AMPure XP beads to the ligation reaction (87uL).
2.8. Follow steps in the AMPure XP bead clean-up section (Appendix E).
2.9. Elute the DNA target from the beads by adding 17 uL of 10mM Tris-HCl or 0.1X TE.
2.10. Transfer 15 uL to a new PCR tube for amplification.

#### 3. PCR enrichment of Adapter-Ligated DNA

3.1. Add the following components to a sterile strip tube:

| Component | Volume (uL) |
|---|---|
| Adapter ligated DNA fragments (from above) | 15 |
| Unique dual index primer pair* | 10 |
| NEBNext Ultra II Q5 master mix (blue tube) | 25 |
| Total volume | 50 |

*The primers are provided in NEBNext® Multiplex Oligos for Illumina® (96 Unique Dual Index Primer Pairs). Please refer to the NEB #E6440 manual for valid barcode combination and tips for setting up PCR reactions

3.2. Set a pipette to 40 uL and then pipette the entire volume up and down at least 10 times to mix thoroughly. Perform a quick spin to collect all liquid from the sides of the tube.
3.3. Place tube on a thermocycler and perform PCR amplification using the following PCR cycling conditions:

| Cycle step | Temperature | Time | # of cycles |
|---|---|---|---|
| Initial denaturation | 98°C | 30 seconds | 1 |
| Denaturation | 98°C | 10 seconds | 3-15* |
| Annealing/extension | 65°C | 75 seconds | |
| Final extension | 65°C | 5 minutes | 1 |
| Hold | 4°C | ∞ | |

*Follow the recommendations for cycle number listed in the table below.

Cycle recommendations

| Input DNA in the end prep reaction | # of cycles required to generate a library yield of: | |
|---|---|---|
| | 100 ng | 1 ug |
| 250 ng | 2-3 | 3-4 |
| 100 ng | 3-4 | 4-5 |
| 50 ng | 4-5 | 5-6 |
| 10 ng | 6-7 | 8-9 |
| 5 ng | 7-8 | 9-10 |
| 1 ng | 8-10 | 11-12 |
| 0.5 ng | 9-10 | 12-13 |
| 0.1ng | 12-13 | N/A |

3.4. Add 0.9X AMPure XP beads to the PCR reactions (45uL).
3.5. Follow steps in the AMPure XP bead clean-up section (Appendix E).
3.6. Elute DNA target from beads into 33 uL 0.1X TE.
3.7. Transfer 30 uL supernatant to a new PCR tube. Libraries can be store at -20C°.
3.8. Check size distribution of libraries and quantitate library concentration.

#### 4. Sizing and quantitation

4.1. Allow TapeStation reagents to equilibrate at room temperature for 30 minutes prior to use.
4.2. Vortex reagents well before use.
4.3. To prepare ladder, mix 2 uL high sensitivity D1000 sample buffer with 2 uL high sensitivity D1000 ladder.

4.4. To prepare sample, mix 2 uL high sensitivity D1000 sample buffer with 2 uL sample.
4.5. Spin down, then vortex using IKA vortexer and adapter at 2000 rpm for 1 minute.
4.6. Spin down to position the sample at the bottom of the tube.
4.7. Load samples into the 2200 TapeStation instrument and follow the software procedure for analysis.
4.8. Quantitate 1 uL library sample with Qubit dsDNA HS kit (Appendix F).

### MiSeq sequencing

#### Protocol Notes

This procedure requires Illumina-style libraries that have been quality-controlled and quantitated using the recommended procedures (i.e. TapeStation and Qubit or qPCR). Exact loading concentrations may vary by machine or lab-dependent factors. For more details on loading and running the MiSeq, consult the more detailed manuals at http://www.illumina.com.

#### Required Reagents

| Company | Product | Catalog number |
|---|---|---|
| Illumina | MiSeq reagent kit v3 | MS-102-3003 |
| Illumina | PhiX control kit v3 | FC-110-3001 |
| | NaOH | |
| | Nuclease-free water | |

#### Procedure

##### 1. Dilute and Pool Libraries

1.1. Calculate the molar concentration of each library to be diluted using average size from the TapeStation and mass from Qubit, using the following equation:

$$\frac{concentration\ (ng/uL)}{660g/mol \times avg\ library\ fragment\ size} \times 10^6\ uL/L = concentration\ (nM)$$

1.2. Make a 4nM dilution of each library.

1.3. Combine equal volumes of each diluted library into a new tube. This is the 4nM library pool.

##### 2. Denature Libraries

2.1. Make a fresh dilution of 0.2N NaOH by combining the following volumes in a microcentrifuge tube:

800 uL laboratory-grade water

200 stock 1.0N NaOH

2.2. Remove HT1 from freezer and thaw at room temperature. Store at 2°C to 8°C until you are ready to dilute denatured libraries.

2.3. Combine the following volumes in a microcentrifuge tube:

5 uL 4nM library

5 uL 0.2N NaOH

2.4. Vortex briefly and then centrifuge at 280 x g for 1 minute.

2.5. Incubate at room temperature for 5 minutes.

2.6. Add 990 uL pre-chilled HT1 to the tube containing denatured library. The result is 1 mL of a 20pM denatured library.

2.7. Dilute the 20pM library to the desired concentration, see table below:

| Concentration | 6 pM | 8 pM | 10 pM | 12 pM | 15 pM | 20 pM |
|---|---|---|---|---|---|---|
| 20 pM library | 180 uL | 240 uL | 300 uL | 360 uL | 450 uL | 600 uL |
| Pre-chilled HT1 | 420 uL | 360 uL | 300 uL | 240 uL | 150 uL | 0 uL |

2.8. Invert to mix and then pulse centrifuge.

2.9. Dilute stock PhiX to 4nM by combining:

2 uL 10 nM PhiX library

3 uL 10 mM Tris-Cl, pH 8.5 with 0.1% Tween 20

2.10. Denature the PhiX control by adding the following volumes in a microcentrifuge tube:

5 uL 4nM PhiX library

5 uL 0.2N NaOH

Remaining 4nM PhiX can be frozen and reused

2.11. Vortex briefly to mix and centrifuge at 280 x g for 1 minute.

2.12. Incubate at room temperature for 5 minutes.

2.13. Dilute denatured PhiX library to 20 pM by adding 990 uL pre-chilled HT1 to the PhiX tube. Invert to mix.

2.14. If using a MiSeq reagent kit v2, dilute 20 pM PhiX library to 12.5 pM by adding the following volumes in a microcentrifuge tube:

375 uL 20pM denatured PhiX library

225 uL pre-chilled HT1

2.15. Combine library and PhiX control according to the table below:

| | |
|---|---|
| Denatured and diluted PhiX | 30 uL |
| Denatured and diluted library | 570 uL |

2.16. Set aside on ice until you are ready to load it onto the reagent cartridge.

#### 3. Load and Run MiSeq

3.1. Thaw frozen reagents overnight at 4°C overnight or in a RT water bath.

3.2. Mix reagents thoroughly by inverting several times. Inspect the bottom of reagent cartridge to ensure all liquids return to the bottom of each tube without any air bubbles.

3.3. Using a 1000 uL pipette tip, piece the foil on position 17.

3.4. Using a fresh 1000 uL pipette tip, transfer the denatured and library (with PhiX spiked) into position 17.

3.5. Generate Sample Sheet using MiSeq Experiment Manager.

3.6. Load MiSeq according to onscreen instructions in the MiSeq Control software.

### 4. Generation of consensus sequences from MiSeq data

The Illumina MiSeq provides very high-quality data, and consensus sequenced may be generated by variety of methods, including commercial tools such as Geneious and CLC Genomics Workbench. The procedure outlined here is a suggestion using free, open source tools.

| Software | Source URL |
|---|---|
| cutadapt 2.3+ | https://github.com/marcelm/cutadapt |
| bowtie2 | https://github.com/BenLangmead/bowtie2 |
| seqtk | https://github.com/lh3/seqtk |
| SAMtools 1.9 | http://www.htslib.org/ |
| BCFtools 1.9 | http://www.htslib.org/ |
| IGV | http://software.broadinstitute.org/software/igv/ |

4.1. Trim reads for quality (Q25+) and for adapters on both ends. Then trim primer sequences (a hard 30 bases on each end), keeping only sequenced that are at least 75 bases. For reads <150 bases, this will need to be modified.

```
cutadapt -j $threads -g GTTTCCCAGTCACGATA -G GTTTCCCAGTCACGATA  \
   -a TATCGTGACTGGGAAAC -A TATCGTGACTGGGAAAC -g ACACTCTTTCCCTACACGACGCTCTTCCGATCT \
   -G ACACTCTTTCCCTACACGACGCTCTTCCGATCT -a AGATCGGAAGAGCACACGTCTGAACTCCAGTCA \
   -A AGATCGGAAGAGCGTCGTGTAGGGAAAGAGTGT -n 3  -m 75   -q 25  \
   --interleaved $read1 $read2 | cutadapt -j $threads --interleaved -m 75 -u 30 \
   -u -30 -U 30 -U -30 -o $read1.trim.fastq -p $read2.trim.fastq -
```

4.2. Map reads to reference sequence

```
bowtie2-build MN908947.fasta MN908947
bowtie2 --sensitive-local  -p $threads -x MN908947 \
   -1 $read1.trim.fastq -2 $read2.trim.fastq -S $samfile
samtools view -b $samfile | samtools sort - -o $bamfile
samtools index $bamfile
```

4.3. Call variants, generate consensus sequence. This will call positions covered by at least 100 reads.

```
samtools mpileup -aa -d 8000 -uf MN908947.fasta $bamfile |\
   bcftools call -Mc |tee -a $vcf | \
   vcfutils.pl vcf2fq -d 100 -D 100000000|\
   seqtk seq -A - | sed  '2~2s/[actg]/N/g'  > $consensusfasta
```

### Appendix A – Singleplex PCR Primers

| Amplicon | 1st round | Sequence | Size | 2nd round | Sequence | Size |
|---|---|---|---|---|---|---|
| PCR1 | 1F_209_1 | GTTGCAGCCGATCATCAGCAC | 756 | W1_2L_368 | TGGAGGAGGTCTTATCAGAGGC | 597 |
| | SC2M1-2_RIGHT_965 | GTTCACGGCAGCAGTATACACC | | SC2M1-2_RIGHT_965 | GTTCACGGCAGCAGTATACACC | |
| PCR2 | W1_2F_00826_1 | AACAACTTCTGTGGCCCTGATG | 904 | W1_2F_00850_2 | TACCCTCTTGAGTGCATTAAAG | 853 |
| | W1_2R_01730_1 | TCCACAAAAGCACTTGTGGAAGC | | W1_2R_01703_2 | AAAGATGCCAAAATAATGGCG | |
| PCR3 | W1_3F_01573_1 | GGTGTTGTTGGAGAAGGTTCCG | 938 | W1_3F_01596_2 | AGGTCTTAATGACAACCTTCTTG | 894 |
| | W1_3R_02511_1 | TGTGGGAAGTGTTTCTCCCTC | | W1_3R_02490_2 | TAAGAAGATAATTTCTTTTGGG | |
| PCR4 | W1_4F_02387_1 | CATTTGTCACGCACTCAAAGGG | 942 | W1_4F_02404_2 | AAGGGATTGTACAGAAAGTGTG | 903 |
| | W1_4R_03329_1 | GTCTGAACAACTGGTGTAAGTTCC | | W1_4R_03307_2 | CCATCTCTAATTGAGGTTGAAC | |
| PCR5 | W1_5F_03185_1 | AGCAAGAAGAAGATTGGTTAGATGATG | 1014 | W1_5F_03208_2 | GATGATAGTCAACAAACTGTTGG | 868 |
| | W1_5R_04199_1 | ATTTCAGTAGTGCCACCAGCC | | W1_5R_04175_2 | TTAGTAGGTATAACCACAGCAG | |
| PCR6 | W1_6F_04054_1 | CATCCAGATTCTGCCACTCTTG | 968 | W1_6F_04073_2 | TTGTTAGTGACATTGACATCAC | 932 |
| | W1_6R_05022_1 | CATGTCCACAACTTGCGTGTG | | W1_6R_05005_2 | TGTGGAGGTTAATGTTGTCTAC | |
| PCR7 | W1_7F_04884_1 | TCCTACCACATTCCACCTAGATGG | 933 | W1_7F_04904_2 | ATGGTGAAGTTATCACCTTTG | 893 |
| | W1_7R_05817_1 | AGCACCGTCTATGCAATACAAAG | | W1_7R_05797_2 | AAGTTTCTTTAGAAGTTATATG | |
| PCR8 | W1_8F_05676_1 | TGTTATGATGTCAGCACCACCTG | 977 | W1_8F_05699_2 | CTCAGTATGAACTTAAGCATGG | 933 |
| | W1_8R_06653_1 | ACAGCAGCTAAACCATGAGTAGC | | W1_8R_06632_2 | GCAAGGGTTTTCAAACCTAATAC | |
| PCR9 | W1_9F_06522_1 | TACAGAAGAGGTTGGCCACAC | 954 | W1_9F_06543_2 | AGATCTAATGGCTGCTTATGTAG | 919 |
| | W1_9R_07476_1 | ACAACCGTCTACAACATGCAC | | W1_9R_07462_2 | CATGCACATAACTTTTCCATAC | |
| PCR10 | W1_10F_07326_1 | TGCAGTACATTTTATTAGTAATTCTTGG | 999 | W1_10F_07356_2 | TATGTGGTTAATAATTAATCTTG | 952 |
| | W1_10R_08325_1 | GTCACGGGGTGTCATGTTTTC | | W1_10R_08308_2 | TTTCAACTTTGTTATAGGTGAGC | |
| PCR11 | W1_11F_08170_1 | GCAGCTCGGCAAGGGTTTGTTG | 952 | W1_11F_08184_2 | GTTTGTTGATTCAGATGTAGAAAC | 922 |
| | W1_11R_09122_1 | CGTGTGTCAGGGCGTAAACTTTC | | W1_11R_09106_2 | AACTTTCATAAGCAACAGAACC | |
| PCR12 | W1_12F_08996_1 | CAGCTTGTGTTTTGGCTGCTG | 980 | W1_12F_09017_2 | AATGTACAATTTTTAAAGATGC | 949 |
| | W1_12R_09976_1 | GAGCCTTTGCGAGATGACAAC | | W1_12R_09966_2 | GAGATGACAACAAGCAGCTTC | |
| PCR13 | W1_13F_09831_1 | GTATCTAAAGTTGCGTAGTGATG | 1005 | W1_13F_09850_2 | GATGTGCTATTACCTCTTACGC | 966 |
| | W1_13R_10836_1 | AACGGCAATTCCAGTTTGAGC | | W1_13R_10816_2 | CAGAAAGAGGTCCTAGTATGTC | |
| PCR14 | W1_14F_10686_1 | TGTTATAAATGGAGACAGGTGG | 984 | W1_14F_10708_2 | TTTCTCAATCGATTTACCACAAC | 949 |
| | W1_14R_11670_1 | GCGGTTGAGTAAACAAAAGAGGC | | W1_14R_11657_2 | CAAAAGAGGCCAAAGTAACAAG | |
| PCR15 | W1_15F_11527_1 | GCCAGAGGTATTGTTTTTATGTGTGT | 911 | W1_15F_11527_2 | GCCAGAGGTATTGTTTTTATGTGTGT | 889 |
| | W1_15R_12438_1 | GGGAACACAACCATCTCTTGC | | W1_15R_12416_2 | TTGTTGATAATGTTGTTGAGTGC | |
| PCR16 | W1_16F_12311_1 | CTAGATCTGAGGACAAGAGGGC | 929 | W1_16F_12327_2 | GAGGGCAAAAGTTACTAGTGC | 892 |
| | W1_16R_13240_1 | ACGATGCACCACCAAAGGATTC | | W1_16R_13219_2 | CTTGATCCATATTGGCTTCCGG | |
| PCR17 | W1_17F_13112_1 | ATCTAGCTAGTGGGGGACAACC | 930 | W1_17F_13126_2 | GGACAACCAATCACTAATTGTG | 902 |
| | W1_17R_14042_1 | AATACCAGCATTTCGCATGGCA | | W1_17R_14028_2 | GCATGGCATCACAGAATTGTAC | |
| PCR18 | W1_18F_13873_1 | TACTTGTCACATACAATTGTTGTGATG | 914 | W1_18F_13873_1 | TACTTGTCACATACAATTGTTGTGATG | 914 |
| | 18R_14809_1 | GATAGTAGTCATAATCGCTGATAGCAG | | W1_18R_14787_1 | TAGCAGCATTACCATCCTGAGC | |
| PCR19 | W1_19F_14655_1 | GCTTTTCAAACTGTCAAACCCGG | 902 | W1_19F_14670_2 | AAACCCGGTAATTTTAACAAAG | 879 |
| | W1_19R_15557_1 | TGCATTAACATTGGCCGTGAC | | W1_19R_15549_2 | CATTGGCCGTGACAGCTTGAC | |
| PCR20 | W1_20F_15429_1 | AGTGAAATGGTCATGTGTGGCG | 971 | W1_20F_15441_2 | ATGTGTGGCGGTTCACTATATG | 939 |
| | W1_20R_16400_1 | ACAACCTGGAGCATTGCAAAC | | W1_20R_16380_2 | CATACGGATTAACAGACAAGAC | |
| PCR21 | 21F_16221_1 | GCATACAGTCTTACAGGCTGTTGG | 919 | W1_21F_16291_2 | GCATACGTAGACCATTCTTATG | 849 |
| | 21R_17140_1 | CAGAAGGGTAGTAGAGAGCTAGGC | | 21R_17140_1 | CAGAAGGGTAGTAGAGAGCTAGGC | |
| PCR22 | W1_22F_17065_1 | ATTCTACACTCCAGGGACCACC | 970 | W1_22F_17082_2 | CCACCTGGTACTGGTAAGAGTC | 930 |
| | W1_22R_18035_1 | TAAAGTTGCCACATTCCTACGTGG | | W1_22R_18012_2 | GAATTTCAAGACTTGTAAATTG | |
| PCR23 | W1_23F_17881_1 | CCACTGAAACAGCTCACTCTTG | 1019 | W1_23F_17901_2 | TGTAATGTAAACAGATTTAATG | 978 |
| | W1_23R_18900_1 | TAACAAAGCACTCGTGGACAGC | | W1_23R_18879_2 | CTAGACACCTAGTCATGATTGC | |
| PCR24 | W1_24F_18767_1 | TGTTCAACAATGGGGTTTTACAGG | 910 | W1_24F_18786_2 | ACAGGTAACCTACAAAGCAACC | 879 |
| | W1_24R_19677_1 | CCTGTTGTCCATCAAAGTGTCCC | | W1_24R_19665_2 | CAAAGTGTCCCTTATTTACAAC | |

| | | | | | | |
|---|---|---|---|---|---|---|
| PCR25 | W1_25F_19546_1 | CAGCTGGCTTTAGCTTGTGGG | 936 | W1_25F_19546_1 | CAGCTGGCTTTAGCTTGTGGG | 936 |
| | 25R_20572_1 | CAACCTTAGAAACTACAGATAAATCTTG | | W1_25R_20482_1 | GATGAACCTGTTTGCGCATCTG | |
| PCR26 | W1_26F_20343_1 | CATAGTCAGTTAGGTGGTTTAC | 972 | W1_26F_20356_2 | GTGGTTTACATCTACTGATTGG | 944 |
| | W1_26R_21315_1 | CTATTTGTTCGCGTGGTTTGCC | | W1_26R_21300_2 | GTTTGCCAAGATAATTACATCC | |
| PCR27 | 27F_21136_1 | AAGCTAGCTCTTGGAGGTTCCG | 926 | W1_27F_21204_2 | CTCATGGGACACTTCGCATGGTGG | 895 |
| | 27R_22218_1 | CCCTGAGGGAGATCACGCAC | | W1_27R_22099_2 | CAAGGTCCATAAGAAAAGGCTG | |
| PCR28 | W1_28F_21976_1 | CCATTTTTGGGTGTTTATTACC | 1017 | W1_28F_21996_2 | CCACAAAAACAACAAAAGTTGG | 979 |
| | W1_28R_22993_1 | TGCTACCGGCCTGATAGATTTC | | W1_28R_22975_2 | TTTCAGTTGAAATATCTCTCTC | |
| PCR29 | W1_29F_22847_1 | TTACAGGCTGCGTTATAGCTTGG | 965 | W1_29F_22864_2 | GCTTGGAATTCTAACAATCTTG | 931 |
| | W1_29R_23812_1 | TGCTGCATTCAGTTGAATCACC | | W1_29R_23795_2 | TCACCACAAATGTACATTGTAC | |
| PCR30 | W1_30F_23681_1 | ACTCTAATAACTCTATTGCCATACCCAC | 944 | W1_30F_23704_2 | CCCACAAATTTTACTATTAGTG | 906 |
| | W1_30R_24625_1 | CAGAAGCTCTGATTTCTGCAGC | | W1_30R_24610_2 | CTGCAGCTCTAATTAATTGTTG | |
| PCR31 | W1_31F_24492_1 | AAATGATATCCTTTCACGTCTTGACAAAG | 999 | W1_31F_24514_2 | GACAAAGTTGAGGCTGAAGTGC | 962 |
| | W1_31R_25491_1 | TTGCAGTAGCGCGAACAAAATC | | W1_31R_25476_2 | CAAAATCTGAAGGAGTAGCATC | |
| PCR32 | W1_32F_25348_1 | CCAGTGCTCAAAGGAGTCAAATTAC | 1019 | W1_32F_25357_2 | AAAGGAGTCAAATTACATTACAC | 1001 |
| | W1_32R_26367_1 | ACGCACACAATCGAAGCGCAG | | W1_32R_26358_2 | ATCGAAGCGCAGTAAGGATGGC | |
| PCR33 | W1_33F_26222_1 | ACAAGCTGATGAGTACGAACTTATG | 906 | W1_33F_26241_2 | CTTATGTACTCATTCGTTTCGG | 874 |
| | W1_33R_27128_1 | TGCCAATCCTGTAGCGACTGTATGC | | W1_33R_27115_2 | GCGACTGTATGCAGCAAAACC | |
| PCR34 | W1_34F_26988_1 | TAGGACGCTGTGACATCAAGG | 1018 | W1_34F_26999_2 | GACATCAAGGACCTGCCTAAAG | 993 |
| | W1_34R_28006_1 | AGGACACGGGTCATCAACTAC | | W1_34R_27992_2 | CAACTACATATGGTTGATGTTG | |
| PCR35 | 35F_27834_1 | ATCTTTTGGTTCTCACTTGAACTGC | 1021 | W1_35F_27875_1 | TGTCACGCCTAAACGAACATG | 980 |
| | 35th_R2_28855 | TGAACTGTTGCGACTACGTGATG | | 35th_R2_28855 | TGAACTGTTGCGACTACGTGATG | |
| PCR36 | W1_36F_28694_1 | CACCAAAAGATCACATTGGCAC | 1030 | W1_36F_28716_2 | CCGCAATCCTGCTAACAATGC | 1008 |
| | W1_36R_29724_1 | TGTGGTGGCTCTTTCAAGTCC | | W1_36R_29724_2 | TGTGGTGGCTCTTTCAAGTCC | |
| PCR37 | W1_37F_29551_1 | AGGCAGATGGGCTATATAAACG | 322 | W1_37F_29596_2 | TATAGTCTACTCTTGTGCAGAATG | 280 |
| | W1_37R_29873_2 | TTTTGTCATTCTCCTAAGAAGC | | W1_37R_29873_2 | TTTTGTCATTCTCCTAAGAAGC | |
| PCR38 | SC2M1-1_LEFT2_1 | TTAAAGGTTTATACCTTCCCAGG | 495 | 0_1b | TTAAAGGTTTATACCTTCCCAGGTA | 490 |
| | SC2M1-1_RIGHT2_495 | CGAGCATCCGAACGTTTGATGA | | W1_1R_490 | CATCCGAACGTTTGATGAACAC | |

### Appendix B – Sequencing Primers

#### Sequencing primer to amplicon matrix

| PCR Product | Sequencing primers | | | |
|---|---|---|---|---|
| PCR1 | W1_2L_368 | SC2M1-2_RIGHT_965 | SC2M1-2_LEFT_445 | SC2M1-1_RIGHT_574 |
| PCR2 | W1_2F_00850_2 | W1_2R_01703_2 | W1_4F_1067* | W1_3R_1206* |
| PCR3 | W1_3F_01596_2 | W1_3R_02490_2 | W1_6L_1819 | W1_5R_1969 |
| PCR4 | W1_4F_02404_2 | W1_4R_03307_2 | W1_9L_2948* | W1_8R_3094* |
| PCR5 | W1_5F_03208_2 | W1_5R_04175_2 | W1_11L_3638* | W1_10R_3792* |
| PCR6 | W1_6F_04073_2 | W1_6R_05005_2 | W1_13L_4307* | W1_12R_4522* |
| PCR7 | W1_7F_04904_2 | W1_7R_05797_2 | W1_15L_5159* | W1_14R_5299* |
| PCR8 | W1_8F_05699_2 | W1_8R_06632_2 | SC2M1-16_LEFT_6030 | SC2M1-15_RIGHT_6172 |
| PCR9 | W1_9F_06543_2 | W1_9R_07462_2 | W1_20L_6877* | W1_19R_7009* |
| PCR10 | W1_10F_07356_2 | W1_10R_08308_2 | W1_22L_7625* | W1_21R_7771* |
| PCR11 | W1_11F_08184_2 | W1_11R_09106_2 | W1_25L_8669* | W1_24R_8794* |
| PCR12 | W1_12F_09017_2 | W1_12R_09966_2 | W1_27L_9308* | W1_26R_9459 |
| PCR13 | W1_13F_09850_2 | W1_13R_10816_2 | W1_29R_10593* | W1_30L_10448* |
| PCR14 | W1_14F_10708_2 | W1_14R_11657_2 | W1_32L_11111* | W1_31R_11251* |
| PCR15 | W1_15F_11527_2 | W1_15R_12416_2 | W1_34L_11808* | W1_33R_11948* |
| PCR16 | W1_16F_12327_2 | W1_16R_13219_2 | W1_37L_12878* | W1_35R_12700* |
| PCR17 | W1_17F_13126_2 | W1_17R_14028_2 | W1_39L_13600* | W1_38R_13741* |
| PCR18 | W1_18F_13873_1 | W1_18R_14787_1 | W1_41L_14342* | W1_40R_14503 |
| PCR19 | W1_19F_14670_2 | W1_19R_15549_2 | W1_43L_14960* | W1_42R_15108* |
| PCR20 | W1_20F_15441_2 | W1_20R_16380_2 | W1_46L_16004* | W1_44R_15773* |
| PCR21 | W1_21F_16291_2 | 21R_17140_1 | W1_48L_16735* | W1_46R_16490* |
| PCR22 | W1_22F_17082_2 | W1_22R_18012_2 | W1_50L_17424* | W1_49R_17553* |
| PCR23 | W1_23F_17901_2 | W1_23R_18879_2 | W1_53L_18503* | W1_52R_18667* |
| PCR24 | W1_24F_18786_2 | W1_24R_19665_2 | W1_55L_19277* | W1_54R_19405* |
| PCR25 | W1_25F_19546_1 | W1_25R_20482_1 | W1_57L_20013* | W1_56R_20146* |
| PCR26 | W1_26F_20356_2 | W1_26R_21300_2 | W1_59L_20656* | W1_58R_20796* |
| PCR27 | W1_27F_21204_2 | W1_27R_22099_2 | W1_61L_21411* | W1_60R_21562 |
| PCR28 | W1_28F_21996_2 | W1_28R_22975_2 | W1_64L_22457 | W1_63R_22612* |
| PCR29 | W1_29F_22864_2 | W1_29R_23795_2 | W1_66L_23182* | W1_65R_23308* |
| PCR30 | W1_30F_23704_2 | W1_30R_24610_2 | W1_69L_24259* | W1_67R_24002* |
| PCR31 | W1_31F_24514_2 | W1_31R_25476_2 | W1_71L_24935* | W1_70R_25075* |
| PCR32 | W1_32F_25357_2 | W1_32R_26358_2 | SC2M1-66_LEFT_25665 | SC2M1-65_RIGHT_25790 |
| PCR33 | W1_33F_26241_2 | W1_33R_27115_2 | SC2M1-68_LEFT_26454 | SC2M1-67_RIGHT_26590 |
| PCR34 | W1_34F_26999_2 | W1_34R_27992_2 | SC2M1-71_LEFT_27650 | SC2M1-69_RIGHT_27432 |
| PCR35 | W1_35F_27875_1 | 35th R2_28855 | W1_81L_28414* | SC2M1-71_RIGHT_28203 |
| PCR36 | W1_36F_28716_2 | W1_36R_29724_2 | SC2M1-75_LEFT_29344 | SC2M1-74_RIGHT_29469 |
| PCR37 | W1_37F_29596_2 | W1_37R_29873_2 | | |
| PCR38 | 0_1b | W1_1R_490 | | |

*Primer sequence in table below.

SC2M1 primers are sourced from the multiplex primer set (Appendix D). Others are primers from Appendix A.

### Additional Sequencing Primer Sequences

| Name | Sequence | Nmae | Sequence |
|---|---|---|---|
| W1_4F_1067 | GGGAATGTCCAAATTTTGTATTTCC | W1_41L_14342 | TTTGGATGACAGATGCATTCTGC |
| W1_3R_1206 | TGGTTGCATTCATTTGGTGACG | W1_43L_14960 | TAAATGGGGTAAGGCTAGACTTTATTATG |
| W1_9L_2948 | TTGATTTAGATGAGTGGAGTATGGCTAC | W1_42R_15108 | CGGTGCGAGCTCTATTCTTTGC |
| W1_8R_3094 | ATGGCTCAAACTCTTCTTCTTCAC | W1_46L_16004 | GATTGAACGGTTCGTGTCTTTAGC |
| W1_11L_3638 | GTGAAGACATTCAACTTCTTAAGAGTGC | W1_44R_15773 | GCTAGCCACTAGACCTTGAGATGC |
| W1_10R_3792 | AGCTAAGTAGACATTTGTGCGAAC | W1_48L_16735 | GGGAAGTTGGTAAACCTAGACCAC |
| W1_13L_4307 | GAACTGTTTCTTGGAATTTGCGAG | W1_46R_16490 | AGCACACAATGGAAAACTAATGGG |
| W1_12R_4522 | CACCATAATCAACCACACCCTC | W1_50L_17424 | GTGTACATTGGCGACCCTGCTC |
| W1_15L_5159 | TTGAGTACTACCACACAACTGATCC | W1_49R_17553 | TTCCGAGGAACATGTCTGGACC |
| W1_14R_5299 | CAGTGGCAAGATAACAGTTGTTATC | W1_53L_18503 | AGGACTTCCTTGGAATGTAGTGCG |
| W1_20L_6877 | AGTGTCGGTAAATTTTGTCTAGAGGC | W1_52R_18667 | CATAGACAACAGGTGCGCTCAG |
| W1_19R_7009 | CAGCGGTTGAGTAGATTAAAGAACC | W1_55L_19277 | TTGTGATGGTGGCAGTTTGTATG |
| W1_22L_7625 | ATTGTGATACATTCTGTGCTGGTAGTAC | W1_54R_19405 | CCATGAGACTCACATGGACTGTC |
| W1_21R_7771 | GATGGATGGAACCATTCTTCACTG | W1_57L_20013 | GACTTATTTAGAAATGCCCGTAATGGTG |
| W1_25L_8669 | TTTCAAGTGAAATCATAGGATACAAGG | W1_56R_20146 | AACTGTGTTTTTACGGCTTCTCC |
| W1_24R_8794 | TACCACCACGCTGGCTAAACC | W1_59L_20656 | AATCTAGTCAAGCGTGGCAACC |
| W1_27L_9308 | CAGGAGTTTTCTGTGGTGTAGATGC | W1_58R_20796 | ATTTTGCGACATTCATCATTATGCC |
| W1_29R_10593 | GTTACCTTCTAAGTCTGTGCCAGC | W1_61L_21411 | CTTAAATTAAGGGGTACTGCTGTTATG |
| W1_30L_10448 | CCAATTTCACTATTAAGGGTTCATTCC | W1_63R_22612 | AAACAGATGCAAATCTGGTGGC |
| W1_32L_11111 | TGGGTATTATTGCTATGTCTGCTTTTG | W1_66L_23182 | TTCAACTTCAATGGTTTAACAGGCAC |
| W1_31R_11251 | TACGCATCACCCAACTAGCAGG | W1_65R_23308 | CAAGTGTCTGTGGATCACGGAC |
| W1_34L_11808 | TGGCAAACCTTGTATCAAAGTAGC | W1_69L_24259 | ATGCAAATGGCTTATAGGTTTAATGG |
| W1_33R_11948 | TGTAACTGGACACATTGAGCCC | W1_67R_24002 | TTGCTTGGTTTTGATGGATCTGG |
| W1_37L_12878 | TCTATACAGAACTGGAACCACCTTG | W1_71L_24935 | ACTGTGATGTTGTAATAGGAATTGTCAAC |
| W1_35R_12700 | AGACATCTGTCGTAGTGCAACAGG | W1_70R_25075 | TGCCAGAGATGTCACCTAAATCAAC |
| W1_39L_13600 | GTCGCTTCCAAGAAAAGGACG | W1_81L_28414 | AATACTGCGTCTTGGTTCACCG |
| W1_38R_13741 | AAGTCATGTTTAGCAACAGCTGG | | |

### Appendix C – Plate Setup for Nested PCR and Sanger Sequencing

Primers are added to each PCR reaction (PCR1-PCR38) prior to adding RNA. The layout stays the same until sequencing reactions are run.

We recommend making PCR primer plates (R1 and R2) in the same format so that primers may be added by multichannel pipetting.

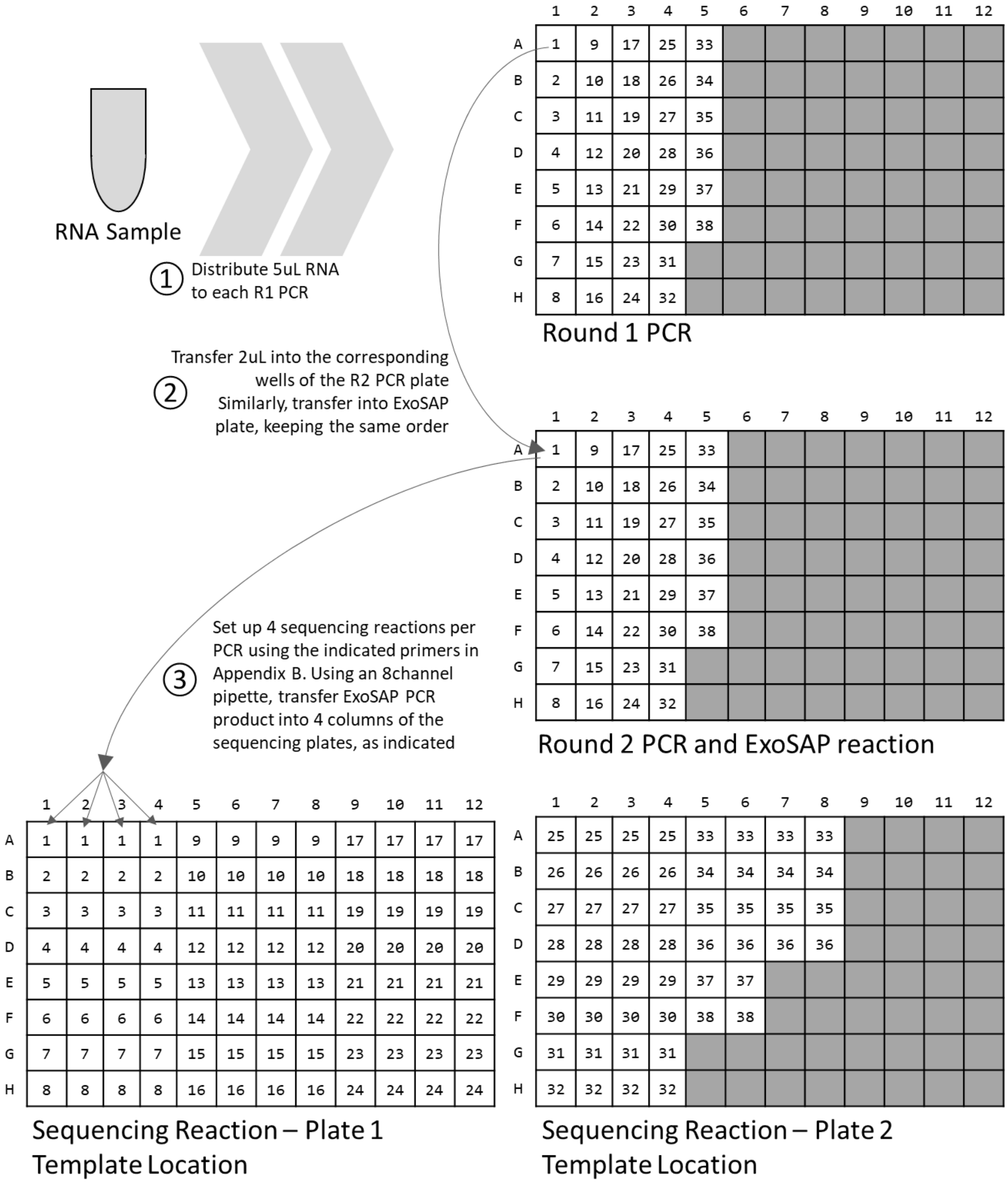

Round 1 PCR

| | 1 | 2 | 3 | 4 | 5 | 6 | 7 | 8 | 9 | 10 | 11 | 12 |
|---|---|---|---|---|---|---|---|---|---|---|---|---|
| A | 1 | 9 | 17 | 25 | 33 | | | | | | | |
| B | 2 | 10 | 18 | 26 | 34 | | | | | | | |
| C | 3 | 11 | 19 | 27 | 35 | | | | | | | |
| D | 4 | 12 | 20 | 28 | 36 | | | | | | | |
| E | 5 | 13 | 21 | 29 | 37 | | | | | | | |
| F | 6 | 14 | 22 | 30 | 38 | | | | | | | |
| G | 7 | 15 | 23 | 31 | | | | | | | | |
| H | 8 | 16 | 24 | 32 | | | | | | | | |

Round 2 PCR and ExoSAP reaction

| | 1 | 2 | 3 | 4 | 5 | 6 | 7 | 8 | 9 | 10 | 11 | 12 |
|---|---|---|---|---|---|---|---|---|---|---|---|---|
| A | 1 | 9 | 17 | 25 | 33 | | | | | | | |
| B | 2 | 10 | 18 | 26 | 34 | | | | | | | |
| C | 3 | 11 | 19 | 27 | 35 | | | | | | | |
| D | 4 | 12 | 20 | 28 | 36 | | | | | | | |
| E | 5 | 13 | 21 | 29 | 37 | | | | | | | |
| F | 6 | 14 | 22 | 30 | 38 | | | | | | | |
| G | 7 | 15 | 23 | 31 | | | | | | | | |
| H | 8 | 16 | 24 | 32 | | | | | | | | |

Sequencing Reaction – Plate 1 Template Location

| | 1 | 2 | 3 | 4 | 5 | 6 | 7 | 8 | 9 | 10 | 11 | 12 |
|---|---|---|---|---|---|---|---|---|---|---|---|---|
| A | 1 | 1 | 1 | 1 | 9 | 9 | 9 | 9 | 17 | 17 | 17 | 17 |
| B | 2 | 2 | 2 | 2 | 10 | 10 | 10 | 10 | 18 | 18 | 18 | 18 |
| C | 3 | 3 | 3 | 3 | 11 | 11 | 11 | 11 | 19 | 19 | 19 | 19 |
| D | 4 | 4 | 4 | 4 | 12 | 12 | 12 | 12 | 20 | 20 | 20 | 20 |
| E | 5 | 5 | 5 | 5 | 13 | 13 | 13 | 13 | 21 | 21 | 21 | 21 |
| F | 6 | 6 | 6 | 6 | 14 | 14 | 14 | 14 | 22 | 22 | 22 | 22 |
| G | 7 | 7 | 7 | 7 | 15 | 15 | 15 | 15 | 23 | 23 | 23 | 23 |
| H | 8 | 8 | 8 | 8 | 16 | 16 | 16 | 16 | 24 | 24 | 24 | 24 |

Sequencing Reaction – Plate 2 Template Location

| | 1 | 2 | 3 | 4 | 5 | 6 | 7 | 8 | 9 | 10 | 11 | 12 |
|---|---|---|---|---|---|---|---|---|---|---|---|---|
| A | 25 | 25 | 25 | 25 | 33 | 33 | 33 | 33 | | | | |
| B | 26 | 26 | 26 | 26 | 34 | 34 | 34 | 34 | | | | |
| C | 27 | 27 | 27 | 27 | 35 | 35 | 35 | 35 | | | | |
| D | 28 | 28 | 28 | 28 | 36 | 36 | 36 | 36 | | | | |
| E | 29 | 29 | 29 | 29 | 37 | 37 | | | | | | |
| F | 30 | 30 | 30 | 30 | 38 | 38 | | | | | | |
| G | 31 | 31 | 31 | 31 | | | | | | | | |
| H | 32 | 32 | 32 | 32 | | | | | | | | |

We recommend making sequencing primer plates as shown below, so that primers may be rapidly added to the sequencing reactions. Primer sequences may be found in Appendices A and D.

| | | | | | | | | | | | |
|---|---|---|---|---|---|---|---|---|---|---|---|
| W1_2L_368 | SC2M1-2_RIGHT_965 | SC2M1-2_LEFT_445 | SC2M1-1_RIGHT_574 | W1_9F_06543_2 | W1_9R_07462_2 | W1_20L_6877 | W1_19R_7009 | W1_17F_13126_2 | W1_17R_14028_2 | W1_39L_13600 | W1_38R_13741 |
| W1_2F_00850_2 | W1_2R_01703_2 | W1_4F_1067 | W1_3R_1206 | W1_10F_07356_2 | W1_10R_08308_2 | W1_22L_7625 | W1_21R_7771 | W1_18F_13873_1 | W1_18R_14787_1 | W1_41L_14342 | W1_40R_14503 |
| W1_3F_01596_2 | W1_3R_02490_2 | W1_6L_1819 | W1_5R_1969 | W1_11F_08184_2 | W1_11R_09106_2 | W1_25L_8669 | W1_24R_8794 | W1_19F_14670_2 | W1_19R_15549_2 | W1_43L_14960 | W1_42R_15108 |
| W1_4F_02404_2 | W1_4R_03307_2 | W1_9L_2948 | W1_8R_3094 | W1_12F_09017_2 | W1_12R_09966_2 | W1_27L_9308 | W1_26R_9459 | W1_20F_15441_2 | W1_20R_16380_2 | W1_46L_16004 | W1_44R_15773 |
| W1_5F_03208_2 | W1_5R_04175_2 | W1_11L_3638 | W1_10R_3792 | W1_13F_09850_2 | W1_13R_10816_2 | W1_29R_10593 | W1_30L_10448 | W1_21F_16291_2 | 21R_17140_1 | W1_48L_16735 | W1_46R_16490 |
| W1_6F_04073_2 | W1_6R_05005_2 | W1_13L_4307 | W1_12R_4522 | W1_14F_10708_2 | W1_14R_11657_2 | W1_32L_11111 | W1_31R_11251 | W1_22F_17082_2 | W1_22R_18012_2 | W1_50L_17424 | W1_49R_17553 |
| W1_7F_04904_2 | W1_7R_05797_2 | W1_15L_5159 | W1_14R_5299 | W1_15F_11527_2 | W1_15R_12416_2 | W1_34L_11808 | W1_33R_11948 | W1_23F_17901_2 | W1_23R_18879_2 | W1_53L_18503 | W1_52R_18667 |
| W1_8F_05699_2 | W1_8R_06632_2 | SC2M1-16_LEFT_6030 | SC2M1-15_RIGHT_6172 | W1_16F_12327_2 | W1_16R_13219_2 | W1_37L_12878 | W1_35R_12700 | W1_24F_18786_2 | W1_24R_19665_2 | W1_55L_19277 | W1_54R_19405 |

Sequencing Reaction – Plate 1
Sequencing Primer Location

| | | | | | | | |
|---|---|---|---|---|---|---|---|
| W1_25F_19546_1 | W1_25R_20482_1 | W1_57L_20013 | W1_56R_20146 | W1_33F_26241_2 | W1_33R_27115_2 | SC2M1-68_LEFT_26454 | SC2M1-67_RIGHT_26590 |
| W1_26F_20356_2 | W1_26R_21300_2 | W1_59L_20656 | W1_58R_20796 | W1_34F_26999_2 | W1_34R_27992_2 | SC2M1-71_LEFT_27650 | SC2M1-69_RIGHT_27432 |
| W1_27F_21204_2 | W1_27R_22099_2 | W1_61L_21411 | W1_60R_21562 | W1_35F_27875_1 | 35th R2_28855 | W1_81L_28414 | SC2M1-71_RIGHT_28203 |
| W1_28F_21996_2 | W1_28R_22975_2 | W1_64L_22457 | W1_63R_22612 | W1_36F_28716_2 | W1_36R_29724_2 | SC2M1-75_LEFT_29344 | SC2M1-74_RIGHT_29469 |
| W1_29F_22864_2 | W1_29R_23795_2 | W1_66L_23182 | W1_65R_23308 | W1_37F_29596_2 | W1_37R_29873_2 | | |
| W1_30F_23704_2 | W1_30R_24610_2 | W1_69L_24259 | W1_67R_24002 | 0_1b | W1_1R_490 | | |
| W1_31F_24514_2 | W1_31R_25476_2 | W1_71L_24935 | W1_70R_25075 | | | | |
| W1_32F_25357_2 | W1_32R_26358_2 | SC2M1-66_LEFT_25665 | SC2M1-65_RIGHT_25790 | | | | |

Sequencing Reaction – Plate 2
Sequencing Primer Location

### Appendix D – Multiplex PCR Primers

#### Pool 1

| PCR | Name | Sequence |
|---|---|---|
| 1 | SC2M1-1_LEFT_31 | ACCAACCAACTTTCGATCTCTTGT |
| | SC2M1-1_RIGHT_574 | TGTCTCACCACTACGACCGTAC |
| 5 | SC2M1-5_LEFT_1706 | TCTGCTTCCACAAGTGCTTTTGT |
| | SC2M1-5_RIGHT_2266 | ACAGGTGACAATTTGTCCACCG |
| 9 | SC2M1-9_LEFT_3306 | TGGAACTTACACCAGTTGTTCAGAC |
| | SC2M1-9_RIGHT_3878 | CAGCGATCTTTTGTTCAACTTGCT |
| 13 | SC2M1-13_LEFT_4885 | TCCTACCACATTCCACCTAGATGG |
| | SC2M1-13_RIGHT_5400 | GCACAAAAGTTAGCAGCTTCACC |
| 17 | SC2M1-17_LEFT_6408 | CTGAAGAAGTAGTGGAAAATCCTACCA |
| | SC2M1-17_RIGHT_6903 | GCCTCTAGACAAAATTTACCGACACT |
| 21 | SC2M1-21_LEFT_8004 | TTGGTGATAGTGCGGAAGTTGC |
| | SC2M1-21_RIGHT_8553 | CCACCCTTAAGTGCTATCTTTGTTGT |
| 25 | SC2M1-25_LEFT_9551 | CCAGTTTACTCATTCTTACCTGGTGT |
| | SC2M1-25_RIGHT_10061 | AACCACTCTGCAAAACAGCTGA |
| 29 | SC2M1-29_LEFT_11047 | AGTCCAGAGTACTCAATGGTCTTTGT |
| | SC2M1-29_RIGHT_11541 | ACAATACCTCTGGCCAAAAACATGA |
| 33 | SC2M1-33_LEFT_12557 | ATCCAACAGGTTGTAGATGCAGAT |
| | SC2M1-33_RIGHT_13136 | TTGGTTGTCCCCCACTAGCTAG |
| 37 | SC2M1-37_LEFT_14103 | TTTCATACAAACCACGCCAGGT |
| | SC2M1-37_RIGHT_14641 | GTGCAGCTACTGAAAAGCACGT |
| 41 | SC2M1-41_LEFT_15637 | AGAAATAGAGATGTTGACACAGACTTTGT |
| | SC2M1-41_RIGHT_16208 | GCCTCATAAAACTCAGGTTCCCA |
| 45 | SC2M1-45_LEFT_17317 | AATGCATTGCCTGAGACGACAG |
| | SC2M1-45_RIGHT_17903 | CAAGAGTGAGCTGTTTCAGTGGT |
| 49 | SC2M1-49_LEFT_18897 | TGTTAAGCGTGTTGACTGGACT |
| | SC2M1-49_RIGHT_19484 | GCACCACCTAAATTGCAACGTG |
| 53 | SC2M1-53_LEFT_20554 | TCTGTAGTTTCTAAGGTTGTCAAAGTGA |
| | SC2M1-53_RIGHT_21144 | AGCTAGCTTTTGTTGTATAAACCCACA |
| 57 | SC2M1-57_LEFT_22203 | GTGATCTCCCTCAGGGTTTTTCG |
| | SC2M1-57_RIGHT_22697 | ACTTAAAAGTGGAAAATGATGCGGAA |
| 61 | SC2M1-61_LEFT_23737 | AATTCTACCAGTGTCTATGACCAAGAC |
| | SC2M1-61_RIGHT_24231 | GCACCAAAGGTCCAACCAGAAG |
| 65 | SC2M1-65_LEFT_25214 | CTAGGTTTTATAGCTGGCTTGATTGC |
| | SC2M1-65_RIGHT_25790 | CATTTCCAGCAAAGCCAAAGCC |
| 69 | SC2M1-69_LEFT_26877 | CTTCTCAACGTGCCACTCCATG |
| | SC2M1-69_RIGHT_27432 | AGCGAGTGTTATCAGTGCCAAG |
| 73 | SC2M1-73_LEFT_28525 | TGGCTACTACCGAAGAGCTACC |
| | SC2M1-73_RIGHT_29045 | GCTTCTTAGAAGCCTCAGCAGC |

#### Pool 2

| PCR | Name | Sequence |
|---|---|---|
| 2 | SC2M1-2_LEFT_445 | TTTGCCTCAACTTGAACAGCCC |
| | SC2M1-2_RIGHT_965 | GTTCACGGCAGCAGTATACACC |
| 6 | SC2M1-6_LEFT_2138 | AAACCCGTCCTTGATTGGCTTG |
| | SC2M1-6_RIGHT_2642 | TTTCGAGCAACATAAGCCCGTT |
| 10 | SC2M1-10_LEFT_3715 | AGCTGGTATTTTTGGTGCTGACC |
| | SC2M1-10_RIGHT_4262 | CCTGACCCGGGTAAGTGGTTAT |
| 14 | SC2M1-14_LEFT_5258 | ACTTCTATTAAATGGGCAGATAACAACTGT |
| | SC2M1-14_RIGHT_5818 | AGCACCGTCTATGCAATACAAAGT |
| 18 | SC2M1-18_LEFT_6748 | AAACCGTGTTTGTACTAATTATATGCCTT |
| | SC2M1-18_RIGHT_7255 | TGCCAAAAACCACTCTGCAACT |
| 22 | SC2M1-22_LEFT_8407 | CGTTAAAGATTTCATGTCATTGTCTGAACA |
| | SC2M1-22_RIGHT_8913 | TGCAAAAAGTCACCATTAGTTGTGC |
| 26 | SC2M1-26_LEFT_9903 | AGTACAAGTATTTTAGTGGAGCAATGGA |
| | SC2M1-26_RIGHT_10451 | TGGGCCTCATAGCACATTGGTA |
| 30 | SC2M1-30_LEFT_11400 | TGAATGTCTTGACACTCGTTTATAAAGTT |
| | SC2M1-30_RIGHT_11944 | CTGGACACATTGAGCCCACAAT |
| 34 | SC2M1-34_LEFT_13006 | TGCCACAGTACGTCTACAAGCT |
| | SC2M1-34_RIGHT_13501 | GTGTAAGACGGGCTGCACTTAC |
| 38 | SC2M1-38_LEFT_14480 | ACTTCAGAGAGCTAGGTGTTGTACA |
| | SC2M1-38_RIGHT_15027 | TGCGAAAAGTGCATCTTGATCCT |
| 42 | SC2M1-42_LEFT_16065 | GGAGTATGCTGATGTCTTTCATTTGTAC |
| | SC2M1-42_RIGHT_16648 | GCGTTTCTGCTGCAAAAAGCTT |
| 46 | SC2M1-46_LEFT_17752 | TGGAGAAAAGCTGTCTTTATTTCACCT |
| | SC2M1-46_RIGHT_18275 | GCTTCTTCGCGGGTGATAAACA |
| 50 | SC2M1-50_LEFT_19311 | TGCATTCCACACACCAGCTTTT |
| | SC2M1-50_RIGHT_19866 | ATTAGCAGCAATGTCCACACCC |
| 54 | SC2M1-54_LEFT_20990 | TGATTGGTGATTGTGCAACTGTACA |
| | SC2M1-54_RIGHT_21562 | TGTTCGTTTAGTTGTTAACAAGAACATCA |
| 58 | SC2M1-58_LEFT_22563 | ACTTGTGCCCTTTTGGTGAAGT |
| | SC2M1-58_RIGHT_23128 | TGCTGGTGCATGTAGAAGTTCA |
| 62 | SC2M1-62_LEFT_24095 | GCTGCTAGAGACCTCATTTGTGC |
| | SC2M1-62_RIGHT_24623 | AAGCTCTGATTTCTGCAGCTCT |
| 66 | SC2M1-66_LEFT_25665 | CTCACACCTTTTGCTCGTTGCT |
| | SC2M1-66_RIGHT_26224 | GTGCTTACAAAGGCACGCTAGT |
| 70 | SC2M1-70_LEFT_27254 | TTATGAGGACTTTTAAAGTTTCCATTTGGA |
| | SC2M1-70_RIGHT_27808 | AGCAGAAAGGCTAAAAAGCACAAA |
| 74 | SC2M1-74_LEFT_28918 | TGATGCTGCTCTTGCTTTGCTG |
| | SC2M1-74_RIGHT_29469 | TCTGCAGCAGGAAGAAGAGTCA |
| 78 | SC2M1-52_LEFT2_20349 | AGTCAGTTAGGTGGTTTACATCTACTGA |
| | SC2M1-52_RIGHT2_20798 | TTTTGCGACATTCATCATTATGCCT |

#### Pool 3

| PCR | Name | Sequence |
|---|---|---|
| 3 | SC2M1-3_LEFT_827 | AACAACTTCTGTGGCCCTGATG |
| | SC2M1-3_RIGHT_1395 | TCTGAATTGTGACATGCTGGACA |
| 11 | SC2M1-11_LEFT_4126 | GGGTGATGTTGTTCAAGAGGGT |
| | SC2M1-11_RIGHT_4658 | ACCGAGCAGCTTCTTCCAAATT |
| 15 | SC2M1-15_LEFT_5677 | TGTTATGATGTCAGCACCACCTG |
| | SC2M1-15_RIGHT_6172 | AGCCACCACATCACCATTTAAGT |
| 19-2 | SC2M1-19b_LEFT_7235 | TGCAGAGTGGTTTTTGGCATATATTCT |
| | SC2M1-19_RIGHT_7694 | ACTGTAGTGACAAGTCTCTCGCA |
| 23 | SC2M1-23_LEFT_8778 | TTAGCCAGCGTGGTGGTAGTTA |
| | SC2M1-23_RIGHT_9330 | TCTACACCACAGAAAACTCCTGGT |
| 27 | SC2M1-27_LEFT_10318 | GCTTAAGGTTGATACAGCCAATCCT |
| | SC2M1-27_RIGHT_10837 | AACGGCAATTCCAGTTTGAGCA |
| 31 | SC2M1-31_LEFT_11810 | GGCAAACCTTGTATCAAAGTAGCC |
| | SC2M1-31_RIGHT_12335 | TTGCCCTCTTGTCCTCAGATCT |
| 35 | SC2M1-35_LEFT_13366 | AAACACAGTCTGTACCGTCTGC |
| | SC2M1-35_RIGHT_13861 | TGTCACAATTACCTTCATCAAAATGCC |
| 39 | SC2M1-39_LEFT_14888 | ACGATGGTGGCTGTATTAATGCT |
| | SC2M1-39_RIGHT_15391 | GGTGTGACAAGCTACAACACGT |
| 43 | SC2M1-43_LEFT_16518 | AAATACATGTGTTGGTAGCGATAATGTT |
| | SC2M1-43_RIGHT_17087 | GGTGGTCCCTGGAGTGTAGAAT |
| 47 | SC2M1-47_LEFT_18148 | GGTTTATGTGTTGACATACCTGGCA |
| | SC2M1-47_RIGHT_18668 | CATAGACAACAGGTGCGCTCAG |
| 51 | SC2M1-51_LEFT_19725 | TGATGGTGTTGATGTAGAATTGTTTGAA |
| | SC2M1-51_RIGHT_20255 | TCAATTTCCATTTGACTCCTGGGT |
| 55 | SC2M1-55_LEFT_21421 | AGGGGTACTGCTGTTATGTCTTTAAA |
| | SC2M1-55_RIGHT_21916 | AAGTAGGGACTGGGTCTTCGAA |
| 59 | SC2M1-59_LEFT_22986 | CCGGTAGCACACCTTGTAATGG |
| | SC2M1-59_RIGHT_23519 | CCCCTATTAAACAGCCTGCACG |
| 63 | SC2M1-63_LEFT_24493 | AAATGATATCCTTTCACGTCTTGACAAA |
| | SC2M1-63_RIGHT_25003 | TGAGTCTAATTCAGGTTGCAAAGGA |
| 67 | SC2M1-67_LEFT_26096 | AAAATTGTTGATGAGCCTGAAGAACA |
| | SC2M1-67_RIGHT_26590 | ACTAGGTTCCATTGTTCAAGGAGC |
| 71 | SC2M1-71_LEFT_27650 | TGTTCATCAGACAAGAGGAAGTTCA |
| | SC2M1-71_RIGHT_28203 | ACGAACAACGCACTACAAGACT |
| 75 | SC2M1-75_LEFT_29344 | TGACGCATACAAAACATTCCCAC |
| | SC2M1-75_RIGHT_29848 | AAAATCACATGGGGATAGCACTACT |

#### Pool 4

| PCR | Name | Sequence |
|---|---|---|
| 4 | SC2M1-4_LEFT_1262 | ACGGGCGATTTTGTTAAAGCCA |
| | SC2M1-4_RIGHT_1840 | TCACCAATATTCCAGGCACCTTT |
| 8 | SC2M1-8_LEFT_2932 | ACTTACACCACTGGGCATTGATT |
| | SC2M1-8_RIGHT_3461 | CTGCAACACCTCCTCCATGTTT |
| 12 | SC2M1-12_LEFT_4519 | TGGTGCTAGATTTTACTTTTACACCAGT |
| | SC2M1-12_RIGHT_5017 | CACAACTTGCGTGTGGAGGTTA |
| 16 | SC2M1-16_LEFT_6030 | ACGCAAGCTTCGATAATTTTAAGTTTGT |
| | SC2M1-16_RIGHT_6544 | TGTGTGGCCAACCTCTTCTGTA |
| 20 | SC2M1-20_LEFT_7560 | GGTCCTTTTATGTCTATGCTAATGGAGG |
| | SC2M1-20_RIGHT_8128 | TGCAAGTTCAGCTTCTGCAGTT |
| 24 | SC2M1-24_LEFT_9203 | GATTCTGAGTACTGTAGGCACGG |
| | SC2M1-24_RIGHT_9734 | AGAACCAATAGAAATGCTTTGTGGAAA |
| 28 | SC2M1-28_LEFT_10697 | GGAGACAGGTGGTTTCTCAATCG |
| | SC2M1-28_RIGHT_11209 | AGCTACAGTGGCAAGAGAAGGT |
| 32 | SC2M1-32_LEFT_12201 | AGTTGAAGAAGTCTTTGAATGTGGCT |
| | SC2M1-32_RIGHT_12719 | TCTGTCGTAGTGCAACAGGACT |
| 36 | SC2M1-36_LEFT_13727 | GCTGTTGCTAAACATGACTTCTTTAAGT |
| | SC2M1-36_RIGHT_14232 | AGGCTTTGTTAAGTCAGTGTCAACA |
| 40 | SC2M1-40_LEFT_15264 | TGTAGAAAACCCTCACCTTATGGG |
| | SC2M1-40_RIGHT_15771 | AGCCACTAGACCTTGAGATGCA |
| 44 | SC2M1-44_LEFT_16948 | CCTACACTAGTGCCACAAGAGC |
| | SC2M1-44_RIGHT_17458 | GTGCAGGTAATTGAGCAGGGTC |
| 48 | SC2M1-48_LEFT_18506 | GACTTCCTTGGAATGTAGTGCGT |
| | SC2M1-48_RIGHT_19038 | ACCAATGTCGTGAAGAACTGGG |
| 52 | SC2M1-52_LEFT_20124 | TGGAGAAGCCGTAAAAACACAGT |
| | SC2M1-52_RIGHT_20698 | GATTAGGCATAGCAACACCCGG |
| 56 | SC2M1-56_LEFT_21775 | TGGGACCAATGGTACTAAGAGGT |
| | SC2M1-56_RIGHT_22345 | ACCAGCTGTCCAACCTGAAGAA |
| 60 | SC2M1-60_LEFT_23379 | ACCAGGTTGCTGTTCTTTATCAGG |
| | SC2M1-60_RIGHT_23876 | CAGCTATTCCAGTTAAAGCACGGT |
| 64 | SC2M1-64_LEFT_24858 | GCACACACTGGTTTGTAACACAA |
| | SC2M1-64_RIGHT_25369 | TTTGACTCCTTTGAGCACTGGC |
| 68 | SC2M1-68_LEFT_26454 | TCCTGATCTTCTGGTCTAAACGAACT |
| | SC2M1-68_RIGHT_27004 | ATGTCACAGCGTCCTAGATGGT |
| 72 | SC2M1-72_LEFT_28066 | TTGAATTGTGCGTGGATGAGGC |
| | SC2M1-72_RIGHT_28649 | TAGCACCATAGGGAAGTCCAGC |
| 76 | SC2M1-1_LEFT2_1 | TTAAAGGTTTATACCTTCCCAGG |
| | SC2M1-1_RIGHT2_495 | CGAGCATCCGAACGTTTGATGA |

#### Pool 5

| PCR | Name | Sequence |
|---|---|---|
| 5W | W1_5L_1457 | GTAAGGGTGGTCGCACTATTGC |
| | W1_5R_1969 | TTGTTATAGCGGCCTTCTGTAAAAC |
| 81* | SC2M1-19a_LEFT_6957 | TGGTTTTTACTATTAAGTGTTTGCCTAGGT |
| | SC2M1-19a_RIGHT_7393 | TCGGGGCCATTTGTACAAGATT |
| 81* | SC2M1-19a_LEFT_6957 | TGGTTTTTACTATTAAGTGTTTGCCTAGGT |
| | SC2M1-19a_RIGHT_7393 | TCGGGGCCATTTGTACAAGATT |
| 83 | SC2M1-21a_LEFT_7984 | AGGCATTAGTGTCTGATGTTGGTG |
| | SC2M1-21a_RIGHT_8384 | TGACTTTTTGCTACCTGCGCAT |
| 28W | W1_28L_9659 | TCACACCTTTAGTACCTTTCTGGATAAC |
| | W1_28R_10207 | GGTTAAGCATGTCTTCAGAGGTGC |
| 85 | SC2M1-34a_LEFT_12994 | GTAGTTTAGCTGCCACAGTACGT |
| | SC2M1-34a_RIGHT_13399 | AACCTTTCCACATACCGCAGAC |
| 42 | SC2M1-42_LEFT_16065 | GGAGTATGCTGATGTCTTTCATTTGTAC |
| | SC2M1-42_RIGHT_16648 | GCGTTTCTGCTGCAAAAAGCTT |
| 89 | SC2M1-49a_LEFT_18711 | CCTGTTGGCATCATTCTATTGGATTT |
| | SC2M1-49a_RIGHT_19112 | GTCACTACAAGGCTGTGCATCA |
| 91 | SC2M1-50a_LEFT_19181 | TGCCTATTTTGGAATTGCAATGTCG |
| | SC2M1-50a_RIGHT_19569 | AAACCCACAAGCTAAAGCCAGC |
| 93 | SC2M1-51a_LEFT_19661 | TTTGATGGACAACAGGGTGAAGT |
| | SC2M1-51a_RIGHT_20098 | GCTTGTTTGGGACCTACAGATGG |
| 60W | W1_60L_21029 | GGATCTCATTATTAGTGATATGTACGACCC |
| | W1_60R_21562 | TTGTTCGTTTAGTTGTTAACAAGAACATC |
| 64W | W1_64L_22457 | CAAAGTGTACGTTGAAATCCTTCACTG |
| | W1_64R_22993 | TGCTACCGGCCTGATAGATTTC |
| 98 | SC2M1-67a_LEFT_25910 | GGCACAACAAGTCCTATTTCTGAAC |
| | SC2M1-67a_RIGHT_26276 | CGTACCTGTCTCTTCCGAAACG |
| 100 | SC2M1-69a_LEFT_26846 | TGTGGTCATTCAATCCAGAAACTAACA |
| | SC2M1-69a_RIGHT_27226 | ACCTGAAAGTCAACGAGATGAAACA |
| 102 | SC2M1-70a_LEFT_27252 | TTATGAGGACTTTTAAAGTTTCCATTTGGA |
| | SC2M1-70a_RIGHT_27644 | AGGTGAAACTGATCTGGCACGT |
| 71 | SC2M1-71_LEFT_27650 | TGTTCATCAGACAAGAGGAAGTTCA |
| | SC2M1-71_RIGHT_28203 | ACGAACAACGCACTACAAGACT |
| 95 | 0_1b | TTAAAGGTTTATACCTTCCCAGGTA |
| | W1_1R_490 | CATCCGAACGTTTGATGAACAC |
| 36 | SC2M1-36_LEFT_13727 | GCTGTTGCTAAACATGACTTCTTTAAGT |
| | SC2M1-36_RIGHT_14232 | AGGCTTTGTTAAGTCAGTGTCAACA |

#### Pool 6

| PCR | Name | Sequence |
|---|---|---|
| 6W | W1_6L_1819 | AGGTGCCTGGAATATTGGTGAAC |
| | W1_6R_2345 | ATGATAGAGTCAGCACACAAAGC |
| 7ab | SC2M1-7a_LEFT_2491 | AGGGAGAAACACTTCCCACAGA |
| | SC2M1-7b_RIGHT_3165 | AGCAGAAGTGGCACCAAATTCC |
| 84 | SC2M1-21b_LEFT_8240 | TCAATCTGACATAGAAGTTACTGGCG |
| | SC2M1-21b_RIGHT_8618 | GCAGCAACAAAAAGGAACACAAGT |
| 26W | W1_26L_8999 | CTTGTGTTTTGGCTGCTGAATG |
| | W1_26R_9459 | CATAAAATAGTAGGCAAGGCATGTTACTAC |
| 40W | W1_40L_13986 | CGCCAAGCTTTGTTAAAAACAGTAC |
| | W1_40R_14503 | TGTACAACACCTAGCTCTCTGAAGTG |
| 86 | SC2M1-34b_LEFT_13245 | CTGTACTGCCGTTGCCACATAG |
| | SC2M1-34b_RIGHT_13620 | CGTCCTTTTCTTGGAAGCGACA |
| 90 | SC2M1-49b_LEFT_18955 | TGCGGCTTGTAGAAAGGTTCAA |
| | SC2M1-49b_RIGHT_19331 | AAAAGCTGGTGTGTGGAATGCA |
| 92 | SC2M1-50b_LEFT_19395 | AGTCTCATGGAAAACAAGTAGTGTCA |
| | SC2M1-50b_RIGHT_19820 | TGGTACTGGTTTAATGTTGCGCT |
| 94 | SC2M1-51b_LEFT_19957 | AACGATTTGTGCACCACTCACT |
| | SC2M1-51b_RIGHT_20373 | TCAGTAGATGTAAACCACCTAACTGACT |
| 99 | SC2M1-67b_LEFT_26128 | TCACACAATCGACGGTTCATCC |
| | SC2M1-67b_RIGHT_26541 | GTACCGTTGGAATCTGCCATGG |
| 101 | SC2M1-69b_LEFT_27080 | TAGCAGGTGACTCAGGTTTTGC |
| | SC2M1-69b_RIGHT_27443 | AAGCTCACAAGTAGCGAGTGTT |
| 9 | SC2M1-9_LEFT_3306 | TGGAACTTACACCAGTTGTTCAGAC |
| | SC2M1-9_RIGHT_3878 | CAGCGATCTTTTGTTCAACTTGCT |
| 75 | SC2M1-75_LEFT_29344 | TGACGCATACAAAACATTCCCAC |
| | SC2M1-75_RIGHT_29848 | AAAATCACATGGGGATAGCACTACT |
| 70b | SC2M1-70b_LEFT_27497 | TCTTCTGGAACATACGAGGGCA |
| | W1_34R_28006_1 | AGGACACGGGTCATCAACTAC |
| 95 | 0_1b | TTAAAGGTTTATACCTTCCCAGGTA |
| | W1_1R_490 | CATCCGAACGTTTGATGAACAC |
| 19-3 | SC2M1-19b_LEFT_7235 | TGCAGAGTGGTTTTTGGCATATATTCT |
| | W1_21R_7771 | GATGGATGGAACCATTCTTCACTG |
| 62-2 | SC2M1-62a1_LEFT_23993 | ACCAAGCAAGAGGTCATTTATTGAAGA |
| | W1_30R_24625_1 | CAGAAGCTCTGATTTCTGCAGC |

*This amplicon is intentionally doubled—add 2 parts of this primer set

### Appendix E – AMPure XP bead clean-up

Bead-based clean-ups are done at several steps throughout the protocols presented. This covers the basic clean-up steps, make sure to check the specific protocol for the ratio of beads to use.

Depending on the number of samples, the AMPure XP bead clean-up takes about 30-40 minutes.

Required reagents for bead-based clean-up

| Company | Product | Catalog number |
|---|---|---|
| Beckman Coulter | Agencourt AMPure XP beads | A63882 |
| | 10mM Tris-HCl pH 8.0 | |

1. Allow AMPure XP beads to warm to room temperature for at least 30 minutes before using.
2. Vortex AMPure XP beads to re-suspend.
3. Add appropriate ratio of re-suspended AMPure XP beads to the ligation reaction. Mix well by pipetting up and down at least 10 times.
4. Incubate for 5 minutes at room temperature.
5. Place the tube/plate on an appropriate magnetic stand to separate the beads from the supernatant. If necessary, quickly spin the sample to collect the liquid from the sides of the tube or plate wells before placing on the magnetic stand.
6. After the solution is clear (about 5 minutes), carefully remove and discard the supernatant. Be careful not to disturb the beads that contain DNA targets (do not discard beads).
7. Add 200 uL of freshly prepared 80% Ethanol to the tube/plate while in the magnetic stand.
8. Incubate at room temperature for 30 seconds, then carefully remove and discard the supernatant.
9. Add another 200 uL of freshly prepared 80% Ethanol to the tube/plate while in the magnetic stand.
10. Incubate at room temperature for 30 seconds, then carefully remove and discard the supernatant.
11. Air dry the beads for 2 minutes while the tube/plate is on the magnetic stand and with the lid(s) open.

    Caution: Do not over dry the beads. This may result in lower recovery of DNA target.
12. Remove the tube/plate from the magnet. Elute the DNA target from the beads by adding appropriate volume of 10mM Tris-HCl or water.
13. Mix well by pipetting up and down or on a vortex mixer. Incubate for 2 minutes at room temperature. If necessary, quickly spin the sample to collect liquid from the sides of the tube or plate wells before placing on the magnetic stand.
14. Place the tube/plate on the magnetic stand.
15. After the solution is clear (about 5 minutes), transfer to a new tube.

### Appendix F – Quantitation using Qubit

Quantitation is done at several various steps throughout the protocols included and this protocol can be used anytime quantitation is indicated.

#### Required reagents

| Company | Product | Catalog number |
|---|---|---|
| Thermo Fisher | dsDNA HS assay kit | Q32854 |
| Thermo Fisher | dsDNA BR assay kit | Q32850 |
| Thermo Fisher | Qubit assay tubes | Q32856 |

Note: depending on the sample, either the high sensitivity (HS) or broad range (BR) kit may be used, the protocols are the same the only difference is the reagents.

Quantitation takes about 10-20 minutes depending on the number of samples.

#### Procedure

1. Set up the required number 0.5 mL Qubit assay tubes for standards and samples. Note: the standards require two tubes.
2. Label tube lids. Do not label the side of the tube as this could interfere with the sample read.
3. Prepare the Qubit working solution by diluting the Qubit dsDNA HS reagent 1:200 in Qubit dsDNA HS buffer. Use a clean plastic tube each time you prepare Qubit working solution. Do not mix the working solution in a glass container.

   The final volume in each tube must be 200 uL. Each standard tube requires 190 uL of Qubit working solution and each sample tube requires anywhere from 180-199 uL. Prepare sufficient Qubit working solution to accommodate all standards and samples.

4. Add 190 uL of Qubit working solution to each of the tubes used for standards.
5. Add 10 uL of each qubit standard to the appropriate tube, mix by vertexing 2-3 seconds.
6. Add Qubit working solution to individual assay tube, mix by vertexing 2-3 seconds.

   Your sample can be anywhere from 1-20 uL. Add a corresponding volume of Qubit working solution to each assay tube: anywhere from 180-199 uL.

7. Add each sample to the assay tubes containing the correct volume of Qubit working solution, then mix by vertexing 2-3 seconds. The final volume in each tube should be 200 uL.
8. Allow all tubes to incubate at room temperature for 2 minutes.
9. Sample concentration can now be measured on the Qubit Fluorometer.

### Appendix G – CENTRI-SEP 96 Protocol

PRINCETON SEPARATIONS

#### CENTRI-SEP 96
#### Protocol

CENTRI•SEP 96 plates must be allowed to equilibrate to room temperature before use. We recommend that the plates be removed from the refrigerator at the same time the sequencing reactions are initiated. This will allow sufficient time for the plates to warm.

1. Remove the adhesive foil from the bottom and then from the top of the CENTRI-SEP 96 plate.
2. Stack the CENTRI-SEP 96 plate on top of a 96-well wash plate and centrifuge at 1500 x g for 2 minutes. Use an external timer and start timing when the rotor has reached the set speed. Discard the liquid in the wash plate. The gel matrix in the wells should appear opaque at this point.
3. Transfer the samples (20 µL or less) to the individual wells in the CENTRI-SEP 96 plate, taking care to place the samples in the centers of the gel beds.
4. Stack the CENTRI-SEP 96 plate on top of a 96-well collection plate and centrifuge at 1500 x g for 2 minutes.
5. Remove the 96-well collection plate containing the cleaned samples and dry in a speed-vac equipped with the appropriate rotor. Alternatively the plate can be sealed for storage.
